# Optimisation of a DNA extraction protocol for improving the bacterial and fungal classification based on Nanopore sequencing

**DOI:** 10.1101/2023.06.21.545968

**Authors:** May Soe Thu, Vorthorn Sawaswong, Prangwalai Chanchaem, Pavit Klomkliew, Barry J. Campbell, Nattiya Hirankarn, Joanne L. Fothergill, Sunchai Payungporn

**Affiliations:** Joint Chulalongkorn University-University of Liverpool Doctoral Program in Biomedical Sciences and Biotechnology, Faculty of Medicine, Chulalongkorn University, Bangkok 10330, Thailand; Department of Infection Biology & Microbiomes, Institute of Infection, Veterinary & Ecological Sciences, University of Liverpool, Liverpool, L69 3GE, UK; Center of Excellence in Immunology and Immune-Mediated Diseases, Department of Microbiology, Faculty of Medicine, Chulalongkorn University, Bangkok 10330, Thailand; Program in Bioinformatics and Computational Biology, Graduate School, Chulalongkorn University, Bangkok 10330, Thailand; Center of Excellence in Systems Microbiology, Chulalongkorn University, Bangkok 10330, Thailand; Department of Clinical Infection, Microbiology and Immunology, Institute of Infection, Veterinary & Ecological Sciences, University of Liverpool, Liverpool, L69 3GE, UK; Department of Biochemistry, Faculty of Medicine, Chulalongkorn University, Bangkok 10330, Thailand

**Author notes:** Corresponding Authors: Joanne L. Fothergill Sunchai Payungporn. E-mail addresses:* May Soe Thu, Vorthorn Sawaswong, Prangwalai Chanchaem, Pavit Klomkliew, Barry J. Campbell, Joanne L. Fothergill, Nattiya Hirankarn, Sunchai Payungporn.

**Keywords:** Bacteria, Fungi, DNA extraction, 16S rRNA, 18S rRNA, Nanopore sequencing

## Abstract

Ribosomal RNA gene amplicon sequencing is commonly used to evaluate microbiome profiles in health and disease and document the impact of interventional treatments. Long-read nanopore sequencing is attractive since it can provide greater classification at the species level. However, optimised protocols to target marker genes for bacterial and fungal profiling are needed. To achieve an increased taxonomic resolution, we developed extraction and long-amplicon PCR-based approaches using Nanopore sequencing. Three sample lysis conditions were applied to a mock microbial community, including known bacterial and fungal species; the 96 MagBead DNA lysis buffer (ML) alone, incorporating bead-beating (MLB) or bead-beating plus MetaPolyzyme enzymatic treatment (MLBE). Profiling of bacterial comparison, MLB had more statistically different bacterial phyla and genera than the others. For fungal profiling, MLB had a significant increase of Ascomycota and a decline of Basidiomycota, subsequently failing to detect *Malassezia* and *Cryptococcus*. Also, the principal coordinates analysis (PCoA) plot by the Bray-Curtis index showed a significant difference among groups for bacterial (*p =* 0.033) and fungal (*p =* 0.012) profiles. Overall, the microbial profiling and diversity analysis revealed that ML and MLBE have more similarity than MLB for both bacteria and fungi, therefore, bead-beating is not recommended for long-read amplicon sequencing. However, ML alone was suggested as an optimal approach considering DNA yield, classification, reagent cost and hands-on time. This could be an initial proof-of-concept study for simultaneous human microbiome and mycobiome studies.

---

High-throughput sequencing (HTS) technologies have undoubtedly had a major impact on genomic research, allowing the study of unculturable microbial communities [1, 2]. This powerful sequencing approach has provided insight into many niches, allowing unrivalled detail into microbiomes [3-7]. Utilising this data, the increased understanding of the importance of microbiota in maintaining human health has contributed to managing healthcare issues through beneficial modification of the microbiome [8-12].

Amplicon sequencing is a typical application of HTS that effectively allows the study of genetic variation from complicated nucleotide mixtures and is considerably more cost-effective than untargeted shotgun metagenomics [13, 14]. A common approach has been targeting conserved genes, such as the 16S nuclear ribosomal RNA (rRNA) gene, to profile complex communities [15]. The 16S rRNA gene has 9 variable regions (V1-V9), useful for determining the bacterial profile to species level within complex biological samples. However, the workflow is highly sensitive, and biases exist at all stages, from initial specimen collection and storage conditions [16, 17] though to microbial DNA extraction [18, 19], DNA sequencing [20] and bioinformatics analysis [21]. Methodological biases can cause significant variances in the observed microbial profiles, resulting in considerable variation between studies [22, 23]. Standardisation of methodologies has therefore been recognised as a significant necessity within industry and regulatory sectors [24]. In this era, sequence-based bacterial investigation of the full 16S rRNA gene (∼1500 bp) combined with HTS has yielded taxonomic resolution at the species and strain level [25, 26]. Long-read sequencing, such as Nanopore sequencing, gives higher resolution in determining members of particular taxa such as *Bifidobacterium* sp. [27, 28].

Less is known about the fungal community within the human microbiome, i.e., the ‘mycobiome’. Fungi can reside within the microbiota, with fungal signatures found in different body sites, including in the buccal cavity, the respiratory, intestinal and urinary tracts, and even breast milk [29]. However, little is known about their interactions with other microorganisms [30], and the fungal profile in the gut accounts for < 1% of the human microbiome [31]. Even so, the global burden of fungal diseases is rising in human immunodeficiency virus (HIV)-infected patients, and infections are also commonly seen in patients with cancer, especially those receiving chemotherapy, and likewise in patients undergoing immunosuppression to support solid organ and stem cell transplantation, and any individuals taking immunomodulatory therapeutics to treat autoimmune and inflammatory diseases [32-35]. Consequently, different sequencing techniques are being used to profile the mycobiome, using markers such as the internal transcribed spacer (ITS) region of the rRNA operon, small ribosomal subunit or 18S rRNA gene, and the large subunit or 28S rRNA gene [30].

Long-read sequencing chemistries allow the study of low-abundance variants and high-heterogeneity samples by amplifying and sequencing full gene lengths, potentially providing species or strain-level resolution [36]. Hence, PacBio and Nanopore-based sequencing have risen in popularity for microbiome studies, and the chemistry advancement for long-read amplicon sequencing using unique molecular identifiers (UMI) has advanced. The mean error rate for the UMI with a read coverage of 15X (Oxford Nanopore Technology-ONT R10.3), 25X (ONT R9.4.1) and 3X (Pacific Biosciences circular consensus sequencing) are 0.0042%, 0.0041% and 0.0007%, respectively [37]. In contrast, Nanopore sequencing has emerged as an appealing and expanding technique for real-time in-field sequencing of environmental and biological microbial samples. This is due to the low cost and portability of long-read nanopore sequencing platforms and the development of quick protocols and analytical pipelines [38, 39].

The importance of a sample processing pipeline for different sample types has previously been identified [18]. Methods to improve DNA yield include increased cell lysis, bead-beating and/or enzymatic methods evaluated within different studies [40, 41]. For short-read sequencing, bead-beating by ceramic or glass beads is a common method for bacterial cell wall lysis, with various protocols identified to increase the recovery of microbial DNA from faecal samples [42]. Protocols with small beads (0.1 mm) have been shown to provide better recovery of bacterial DNA, while methods with bigger beads (0.5 mm) yielded higher fungal DNA recovery [18, 42], demonstrating that optimal processing to cover all microbial species within samples extensively can be challenging. A combination of bead beating and enzymatic lysis (e.g., lysozyme) is also encouraged, providing both mechanical disruption and enzymatic degradation of bacterial cell wall peptidoglycans, particularly for Gram-positive bacteria, maximising the release of DNA and proteins [43, 44]. However, an optimised, simultaneous extraction protocol for both bacterial and fungal communities for long-read sequencing has yet to be assessed.

In this study, we evaluated three lysis approaches using a commercially available DNA extraction kit and used a reference mock microbiota community containing representative bacterial and fungal species. These were subjected to a long-read Oxford Nanopore sequencing to establish the best method for human gut microbiome studies.

## MATERIALS AND METHODS

### Experimental design

For this study, we generated a mock microbiota community suspension using two commercially available reference standards: the first was the ZymoBIOMICS™ faecal reference kit with TruMatrix™ technology (D6323; Zymo Research, Irvine CA, USA); DNA content, 6 ng/µL comprising of 71% Firmicutes, 23% Bacteroidetes, 1% Actinobacteria, 1% Verrucomicrobia and < 1% for other phyla. The other, the ATCC^®^ MSA-2010^TM^ mycobiome whole cell mix (American Type Culture Collection, Manassas VA, USA); NGS standards; specification range, 2 × 10^7^ cells/vial (± 1 log), comprising of 10 fungal species, each with 10% of the total. Mock microbiota community samples were prepared from equal cell suspension volumes of the faecal reference kit (100 μL) and the mycobiome whole cell mix (100 μL), with duplicates assigned to each of three extraction conditions: 1) use of the 96-MagBead DNA extraction kit lysis buffer alone (ML); 2) lysis buffer with bead-beating (MLB); and 3) lysis buffer, bead-beating and an enzymatic treatment step using MetaPolyzyme (MAC4L; Sigma-Aldrich, St Louis MO, USA) (MLBE). A negative control, with no starting microbial reference suspension, was included. Post-treatment, cellular DNA was extracted, purified and PCR amplified, and then processed for sequencing (Figure 1).

**Figure 1:**
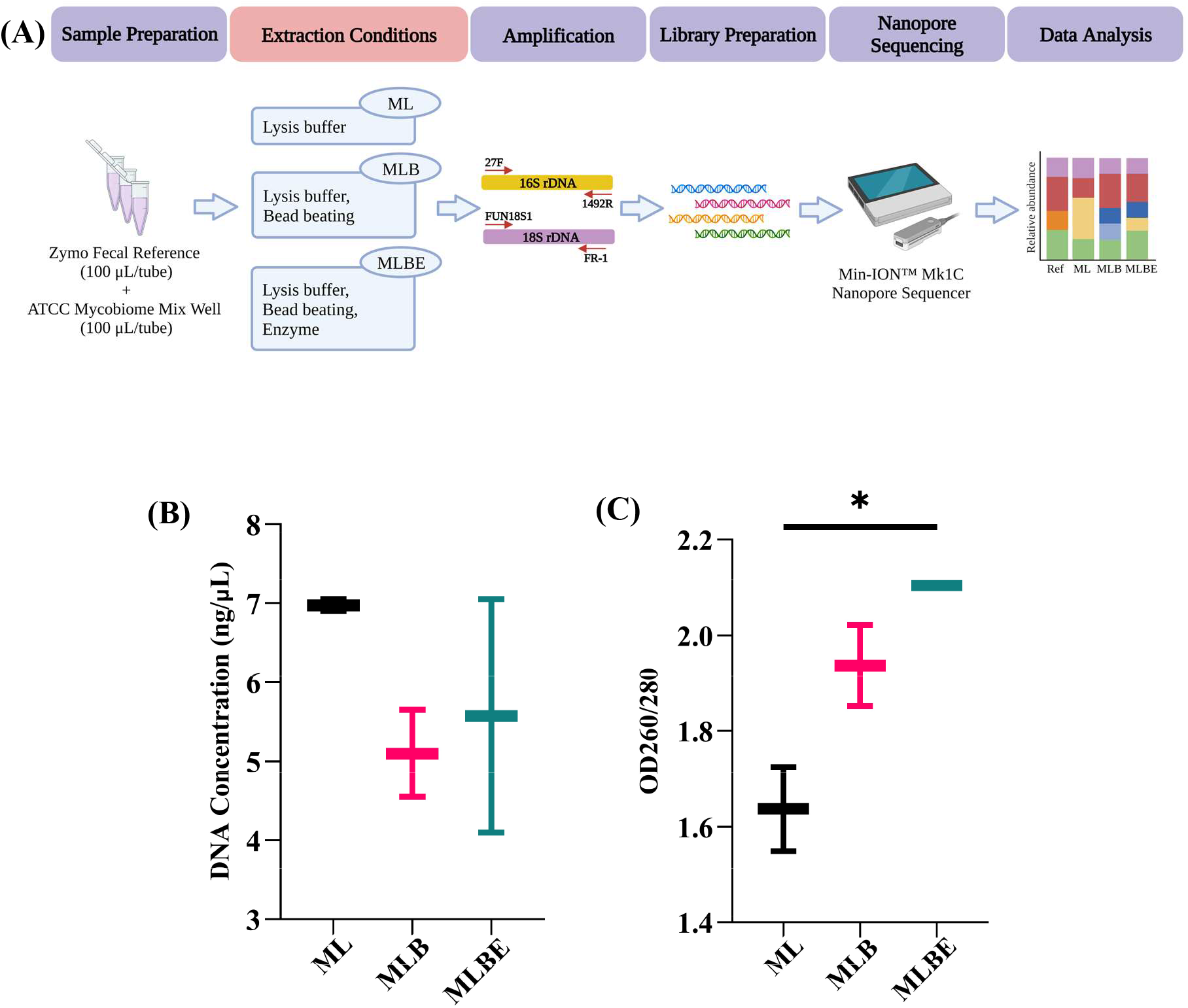
(A) Schematic workflow of optimising different DNA extraction methods for microbiome study using Nanopore long-read sequencing. DNA extraction conditions were as follows; 96 MagBead DNA lysis buffer alone (ML), incorporating bead-beating (MLB), or bead-beating plus MetaPolyzyme enzymatic treatment (MLBE). Following extraction, (B) yield and (C) purity of isolated DNA was assessed by Implen Nanophotometer. Significant differences were observed, \**p* < 0.05 (One-way ANOVA).

### Sample treatments and DNA extraction

All mock microbiota community suspension samples were pre-heated to 95°C and mixed at 900 rpm for 10 min using a Thermomixer^®^ (Eppendorf, Hamburg, Germany). For ML, 750 μL kit lysis solution was added to each cell suspension sample. For MLB, suspensions were transferred into ZR Bashingbead^TM^ lysis tubes containing a mix of 0.1 mm and 0.5 mm zirconium beads (Zymo Research) and 750 μL lysis buffer. For the MLBE aliquots, 750 μL lysis solution and 5 μL MetaPolyzyme were added to cell suspensions. The MLB and MLBE tubes were then subjected to bead-beating for 6 min using the TissueLyser Lt (Qiagen^®^, Hilden, Germany), with the instrument set at 50 Hz, with 1 min rest on ice after every 1.5 min of the bead-beating.

All samples were centrifuged at ≥10,000 x g for 1 min after treatments. For DNA purification, 200 μL of supernatant was added to 600 μL ZymoBIOMICS™ MagBinding buffer and 25 μL of ZymoBIOMICS™ MagBinding beads, mixed well on a shaker plate at 600 rpm for 10 min, then followed as per manufacturer instructions. Quantity and quality of eluted DNA concentration were measured in a NanoPhotometer^®^ C40 machine (Implen; Westlake Village CA, USA), with the OD_260/280_ ratio used to determine the sample purity was within the desired range (1.55 – 2.10). Samples were then stored at -20°C before amplification and sequencing. Samples were stored frozen for less than 1 month prior to extraction.

### Amplification before nanopore sequencing

For bacterial microbiome detection, amplification of the full length of the 16S rRNA gene was performed using forward primer 27F and reverse primer 1492R (Macrogen, Seoul, South Korea) with anchor sequences [28]; for oligonucleotide primer sequences, Table S1. All the amplification was duplicated for sequencing. An in-house developed control of full-length V1-V9 competent *Escherichia coli* (Bangkok, Thailand) was used as a positive control for PCR amplification, and sterile OmniPur diethylpyrocarbonate (DEPC)-treated water (Sigma-Aldrich) was used as a template for the negative control.

For the 2-step PCR, the first PCR reaction comprised 10 ng of DNA template, 1X Phusion^TM^ Plus buffer (F630XL; ThermoFisher Scientific, CA, US), 200 µM dNTPs, 0.2 µM of each primer, 0.5U PCR of Phusion^TM^ Plus DNA polymerase (ThermoFisher Scientific), and DEPC water in a total volume of 20 µL. The PCR was performed on an Applied Biosystems ProFlex™ PCR System (ThermoFisher Scientific) using the following program: 98°C for 30 s for initial denaturation, 25 cycles of 98°C for 10 s, 60°C for 10 s, and 72°C for 45 s for denaturation, annealing and extension steps respectively, and then a final extension of 72°C for 5 min. This allowed amplicon production with overhung adapter sequences, facilitating the second round of PCR. The master mix preparation for the second PCR was as above but using half volume of barcodes and only 5 thermal cycles. This allowed for the addition of unique barcodes to each sample, Table S2.

For the fungal mycobiome amplification, the primer set nu-SSU-0068-5′-20 (Fun18S1) and nu-SSU-1648-3′ (FR-1) (Macrogen, South Korea), targeting the full length (1.6 kbp) 18S rRNA [45] was used, Table S1. The PCR reaction was identical to that performed for 16S rRNA but over 35 thermal cycles [46]. A non-template negative control was also included.

Amplified products were assessed for quality using 2% w/v agarose gel electrophoresis in 1X Tris/Borate/EDTA (TBE) buffer and purified using a QIAquick PCR purification kit (Qiagen). The purified libraries were pooled equimolarly and subsequently purified using 0.5X Agencourt AMPure XP beads (A63882; Beckman Coulter Life Sciences, Indianapolis, IN, USA). Before sequencing, libraries were quantified using a Qubit™ 4 fluorometer and Qubit dsDNA HS (high sensitivity) assay kit (Q33239; ThermoFisher Scientific).

### Nanopore Sequencing

Generated libraries underwent DNA repair, end-prep, adapter ligation and clean-up, priming and loading to the SpotON flow cell according to the manufacturer-recommended Ligation sequencing amplicons protocol (SQK-LSK112, ONT, Oxford, UK). The libraries were loaded onto a Min-ION™ flow cell (vR10.4; FLO-MIN112; ONT), and sequencing was performed on the Min-ION™ Mk1C nanopore sequencer (ONT). Min-ION™ MINKNOW software v5.3.6 (ONT); https://nanoporetech.com/about-us/news/introducing-new-minknow-app; accessed 29 September 2022, was utilised data acquisition.

### Data processing and bioinformatic analysis

To generate the FASTQ files, the super-accuracy model of Guppy basecaller v6.0.1 (ONT) was used to basecall the FAST5 data. The read quality was evaluated using MinIONQC [47]. Then, FASTQ sequences were demultiplexed and adaptors trimmed using Porechop v0.2.4 (https://github.com/rrwick/Porechop; accessed 05 April 2022). The filtered reads were clustered, polished, and taxonomically classified by NanoCLUST [48]. Bacterial taxonomy was classified using RDP database v11.5, and fungal taxonomy by an in-house curated database having 10 fungal species of the mock communities (https://gofile-37314c4275.sg3.quickconnect.to/fsdownload/wbqHvlnID/18S%20project%20(May); accessed 05 April 2022). The abundance taxonomic assignment data were converted into QIIME data format using the QIIME2 platform (https://qiime2.org/; accessed 05 April 2022). The normalised data files were imported into the platform of MicrobiomeAnalyst (www.microbiomeanalyst.ca/; accessed 05 April 2022) to assess microbial richness and evenness based on the relative abundance of taxa, α-diversity (Chao1 and Shannon indexes) and β-diversity (Bray-Curtis index) [49].

### *In Silico* PCR oligonucleotide primer sequence verification

To evaluate the amplification performance of the primer set for the 18S rRNA gene and to determine whether we could detect all 10 fungal species within the mock microbiota community, BioEdit v7.2.5 was utilised. After collecting the full ribosomal sequences of the 10 fungi, CLUSTALW Multiple Sequence Alignment (https://www.genome.jp/tools-bin/clustalw; accessed 13 June 2022) was used to align with the forward and reverse primer set primer sequences used within this study; Figure S1. Multiple alignments of 18S rRNA gene sequences were conducted using the contigs from ATCC genome assembly; ATCC36031 (*Fusarium keratoplasticum*), and the following accession number sequences; CP046435.1 (*Malassezia globosa*), D12804.1 (*Cryptococcus neoformans*), JAADCK010000556.1 (*Trichophyton interdigitale*), M55628.1 (*Penicillium chrysogenum),* M60300.1 (*Aspergillus fumigatus*), NG_062025.1 (*Cutaneotrichosporon dermatis*), X51831.1 (*Candida glabrata*), X53497.1 (*Candida albicans*), and Z75578.1 (*Saccharomyces cerevisiae*).

### Amplification of the genus *Malassezia*

The extracted DNA from different lysis conditions was amplified using *Malassezia*-specific PCR primers: MAL1F (5’-TCTTTGAACGCACCTTGC-3’) and MAL1R (5’-AHAGCAAATGACGTATCATG-3’) (Macrogen, South Korea) which provided ∼300 bp fragment containing 5.8S rRNA gene and ITS2 [50, 51]. PCR was performed using OnePCR Ultra^TM^ Master Mix (MB208-0100; Bio-Helix, New Taipei, Taiwan), 10 pmol of each primer, and 1 µL of extracted DNA in a final 20 µL volume. The PCR was performed on a ProFlex™ PCR System with the program: 94°C for 2 min as an initial denaturation, 40 cycles of 94°C for 20 s, 60°C for 30 s, and 72°C for 2 min as denaturation, annealing, and extension respectively, and then 72°C for 5 min for a final extension. Negative control was also included; Figure S2. The purified amplicons were confirmed by Sanger sequencing on an ABI 3730XL DNA analyser (U2Bio Co. Ltd; Bangkok, Thailand).

### Spike-in Controls

To evaluate the binding efficiency of the primer set (Fun18S1/FR-1) to the *M. globosa* and other fungal species in a matrix of microbial samples, the spike-in experiment was conducted at which control samples were prepared using purified DNA isolated from spores of cultured *M. globosa* (Supplementary Material; Methods S1). We used 50%, 25%, 10%, 5% and 0% spike-in controls by mixing the same DNA concentration (2.35 ng/µL) of combined mock controls and known *M. globosa*, followed by PCR amplification; Figure S2 and Nanopore sequencing; Figure S3.

### Statistical analyses

Comparison of DNA yields achieved using the three lysis methods was conducted by one-way analysis of variance (ANOVA). To determine the significant difference between groups on the abundance of bacterial and fungal profiles, a non-parametric Kruskal-Wallis test was used. Dunn’s multiple pairwise comparison test was applied to analyse differences in the abundance of each sample group to the reference, and differences were considered statistically significant when *p* < 0.05. Shannon’s diversity index was used to determine evenness, and the chao1 index for the richness of operational taxonomic units (OTUs) to identify the α-diversity. A Bray-Curtis test was used for the detection of β-diversity, followed by permutational multivariate analysis of variance using distance matrices (PERMANOVA) test, and visualised using a Principal component analysis (PCoA) plot. Statistical analyses and visualisation were performed using MicrobiomeAnalyst (www.microbiomeanalyst.ca/; accessed 26 June 2022) and GraphPad Prism v9.0 (GraphPad Software, Inc.; La Jolla CA, USA).

## RESULTS

### Average DNA concentration across all extraction methods was not significantly different

To ascertain the basic characteristics of the DNA following extraction, isolated DNA levels from the mock microbial communities were determined and found to be in the range of 4.1-7.05 ng/µL. The average values of DNA concentration in each lysis method were not different (*p* = 0.4264), and DNA purity was within the desired range (OD_260/280_ 1.55 – 2.10); see Figure 1B. However, intra-sample variation in DNA concentration was noted, with the second sample consistently displaying a lower concentration. Further experimentation was performed on the initial sample only.

### Primer sets were optimised using different thermal cycles

PCR amplification is a key step in this approach. The new primer set of 16S rRNA gene was utilised with Phusion plus PCR protocol as described in a previous study [52]. To ensure the DNA quality, we first set up different thermal cycles, and then DNA bands with good intensity were achieved by 25X cycles, which is within the range of the manufacturer’s recommendation. The same PCR condition was used for the 18S rRNA gene amplification. The DNA quality was optimised at 35X cycles for the initial amplification as in the previous research study [53].

### Concentration of DNA libraries and Sequencing reads

For the 16S rRNA gene amplification, the DNA concentration for a couple of samples (ML1_2 and MLBE1_2) was around 10 ng/µL, and one duplicate from each treatment (ML1_1, MLB1_1, MLBE1_1) was >20 ng/μL. The range of the average DNA concentration for the 18S rRNA gene amplification was 1.45-7.20 ng/µL (Table 1). One sample (MLB1_2) yielded a low DNA concentration for both the 16S amplification (4.98 ng/µL) and the 18S amplification (0.638 ng/µL), along with very few sequencing reads (Table 1).

**Table 1:** Concentrations of DNA libraries and numbers of sequencing reads

| Conditions | 16S rRNA |  | 18S rRNA |  |
| --- | --- | --- | --- | --- |
| | Concentration (ng/μL) | Reads | Concentration (ng/μL) | Reads |
| ML1 | 19.00 | 12,034 | 7.20 | 9,755 |
| MLB1 | 15.29 | 13,440 | 1.45 | 3,727 |
| MLBE1 | 19.15 | 14,354 | 7.00 | 7,394 |

### Subsampling

To increase the accuracy of taxonomic profile analysis and make an identical analysis on each sample, a subsampling of 10,000 read counts for the 16S rRNA gene was undertaken. Despite the low concentration at MLB1_2, all samples were taken forward for DNA sequencing to understand the library quality impact on sequencing output. With very low reads on MLB1_2, there was no subsampling for the 18S rRNA gene. The rarefaction plots for both genes were created by MicrobiomeAnalyst; Figure S4.

### Bacterial taxonomic profile using 16S rRNA gene sequencing

At the phylum level, there are three phyla (Firmicutes, Bacteroidetes, and Proteobacteria) within the reference data, while the mock samples have not only these three phyla but also the others, such as Verrucomicrobia and Actinobacteria in all the conditions, and Lentisphaerae only in MLBE1_2 (Figure 2A). Regarding genera, 31 were identified within the reference library, and 46 genera in all samples. For data visualisation, the relative abundance of identified genera was ranked, and the top 15 within the reference data (>70% of the total identified) were presented, with all remaining identified genera (<30%) grouped and represented as ‘Others’ (Figure S5A). Only *Bacteroides vulgatus* and *Bacteroides dorei* were matched to the reference library at the species level, and all others were unknown species (Figure 2B).

**Figure 2:**
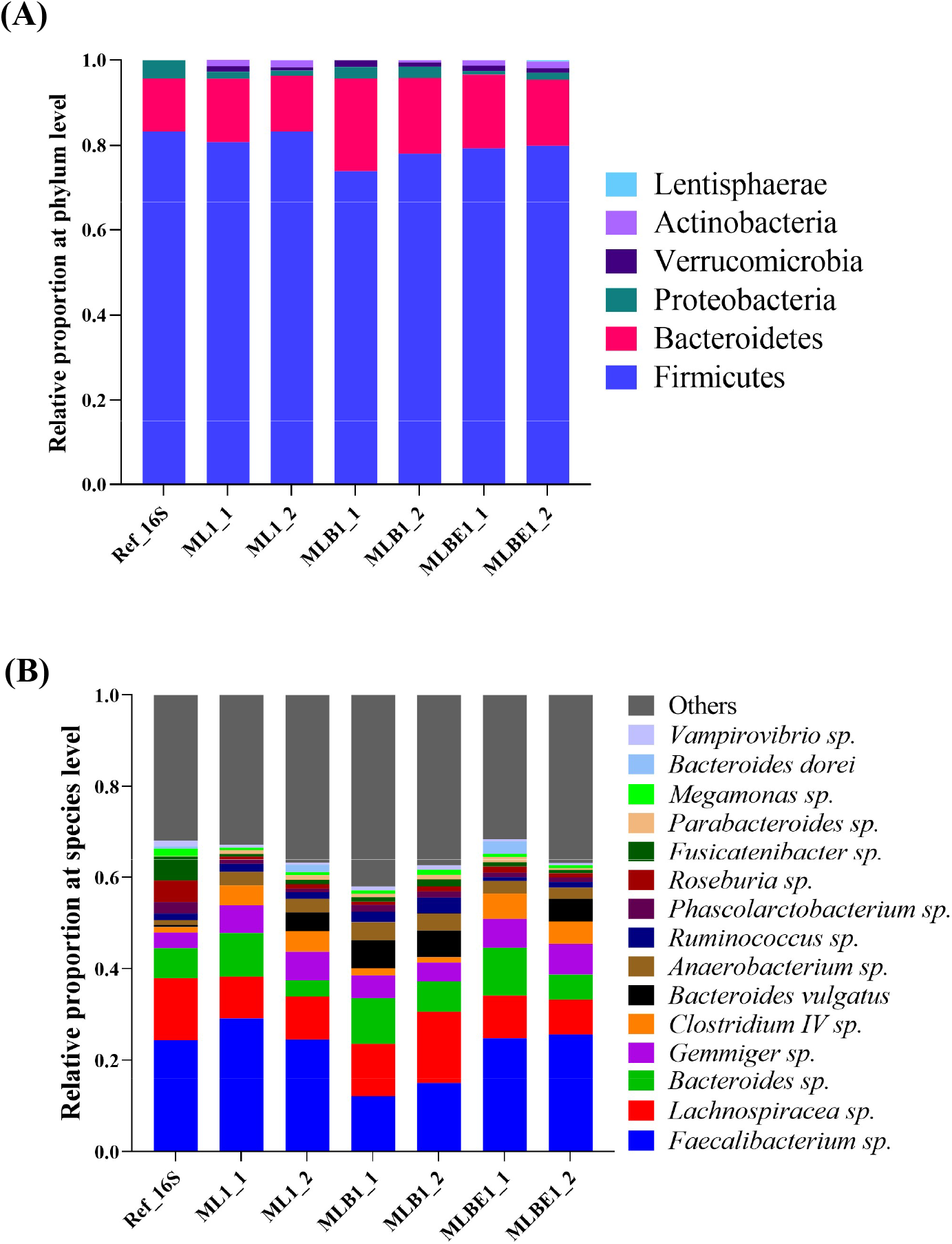
Relative bacterial abundance identified by Nanopore long-read sequencing using DNA isolated by 3 different extraction methods, lysis buffer alone (ML), incorporating bead-beating (MLB), or bead-beating plus MetaPolyzyme enzymatic treatment (MLBE). Data illustrated shows (A) all phyla, and (B) the top 15 species identified. The initial samples: ML1, MLB1, and MLBE1 were duplicated for Nanopore sequencing, abbreviated as ML1_1, ML1_2, MLB1_1, MLB1_2, MLBE1_1 and MLBE1_2, respectively. The reference data of 16S gene sequencing was abbreviated as Ref_16S.

Looking through the individual phyla (Figure 3A-C), no significant differences were found comparing the DNA extraction conditions ML and MLBE; however, a significant increase of Bacteroidetes (Figure 3B) was noted using MLB (*p =* 0.0429, Table S3). We identified 3 common genera *Faecalibacterium*, *Bacteroides* and *Lachnospiracea*, which comprised half of the total bacterial proportions; Figure S6. MLB yielded decreased levels of *Faecalibacterium* (*p* > 0.05, Table S3) and an increase in *Bacteroides* (*p =* 0.0412, Table S3). The abundance of *Lachnospiracea* was seen to be consistent across all extraction groups. Other significant reductions in the relative abundance of genera noted included *Phascolarctobacterium* in the ML extraction conditions (*p =* 0.0412, Table S3) and *Gemmiger* using MLBE (*p =* 0.0412, Table S3), Figure S6. However, these represent a very low proportion of the reference community. No significant differences were found at the species level, Figure S7.

**Figure 3:**
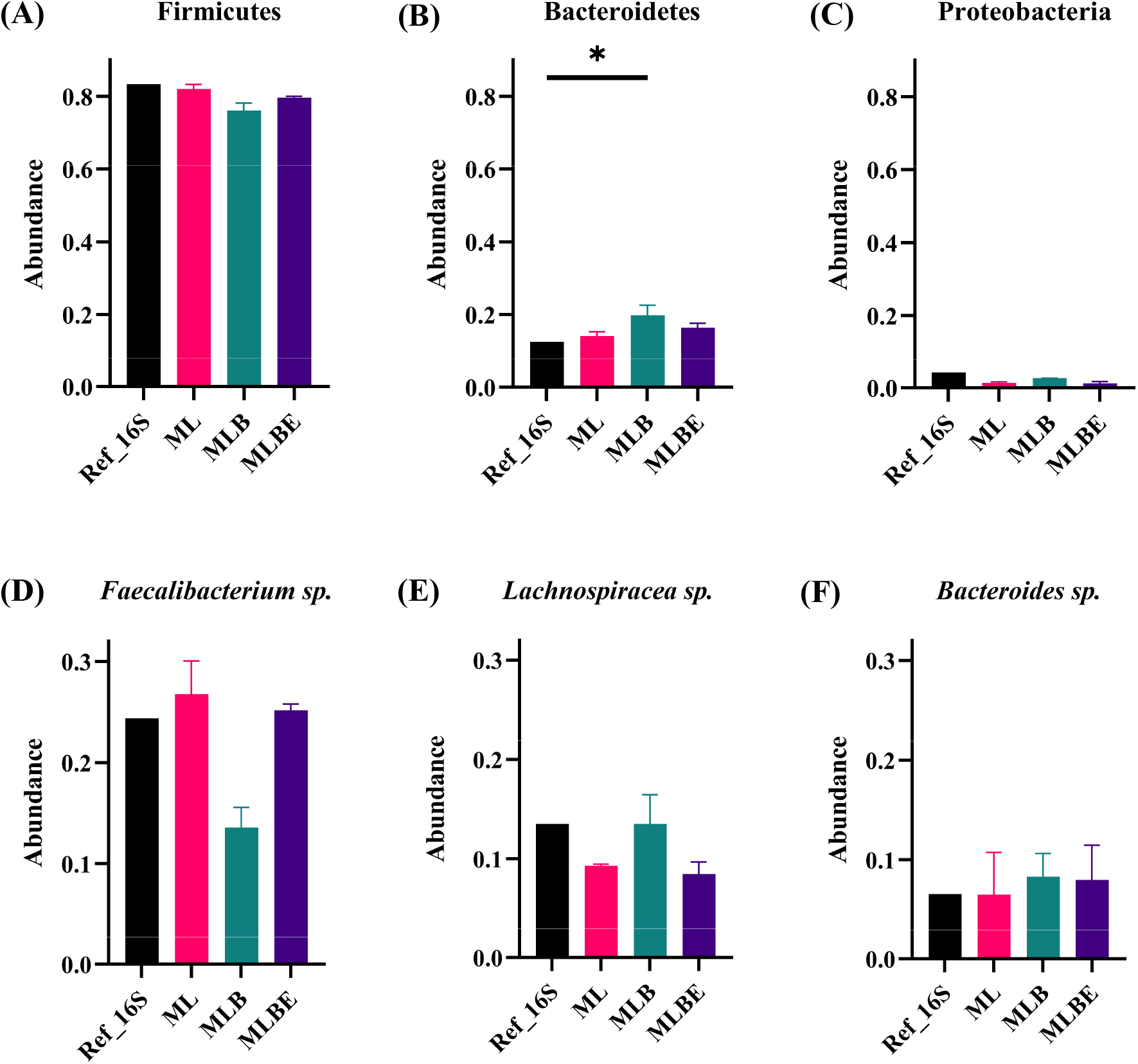
Relative abundance of the top 3 identified bacterial taxa identified by Nanopore sequencing of DNA isolated under different DNA extraction conditions; 96 MagBead DNA lysis buffer alone (ML), incorporating bead-beating (MLB), or bead-beating plus MetaPolyzyme enzymatic treatment (MLBE). Abundance at phylum level; (A) Firmicutes, (B) Bacteroidetes and (C) Proteobacteria. At the species level, (D) *Faecalibacterium* sp.; (E) *Lachnospiracea* sp.; (F) *Bacteroides* sp. Data are expressed as the average of duplicates, and lines indicate median values. Significant differences were observed, \**p* < 0.05 (Kruskal-Wallis test and Dunn’s multiple comparisons posthoc test).

Assessment of β-diversity revealed a significant difference observed between the extraction groups (Figure 4A-B), with treatments ML and MLBE displaying the most similarity to the reference and each other. Conversely, there were no significant differences in α-diversity (Figure 4C-D).

**Figure 4:**
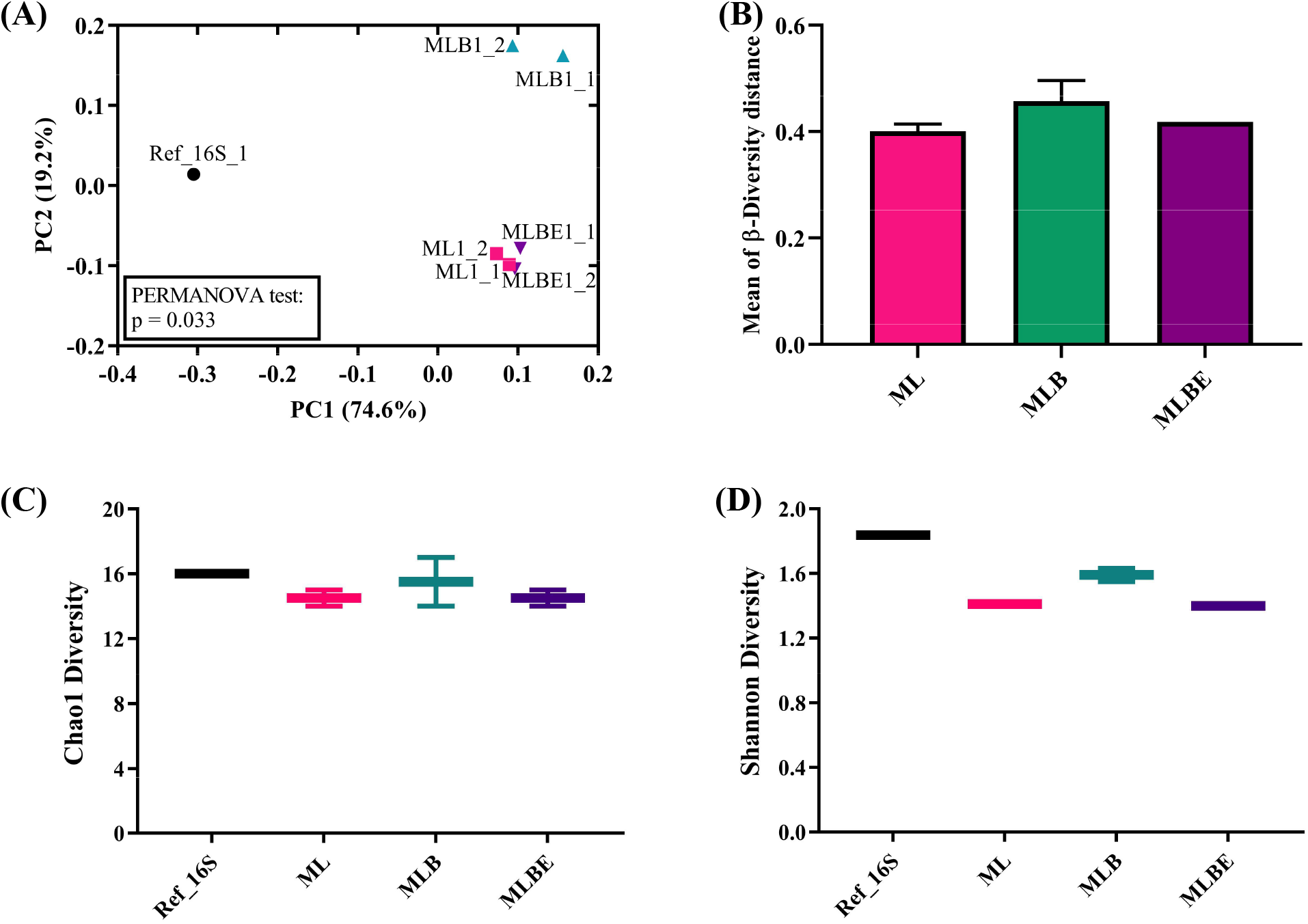
Bacterial diversity in different extraction methods, 96 MagBead DNA lysis buffer alone (ML), incorporating bead-beating (MLB), or bead-beating plus MetaPolyzyme enzymatic treatment (MLBE). Graphs illustrate (A) β-diversity at the species level by Bray-Curtis metrics, (B) the Coordinate distance of the Bray-Curtis index at the bacterial species level, (C) α-diversity at the species level using the Chao1 index, and D) α-diversity at species level using Shannon index. Histogram bars represent the mean ± standard deviation. The line in the data plots indicates the median value. Significant differences were observed, \**p* < 0.05 (Kruskal-Wallis test and Dunn’s multiple comparisons posthoc test).

### Fungal taxonomic profile using 18S rRNA gene sequencing

Using the 18S rRNA gene, the phyla of Basidiomycota and Ascomycota were identified in all samples, but the relative proportions differed dramatically from the reference data (Figure 5A). The mycobiome reference kit has 10 fungal species from 9 genera. However, only 9 species were detected in the samples, with *M. globosa* notably absent (Figure 5B).

**Figure 5:**
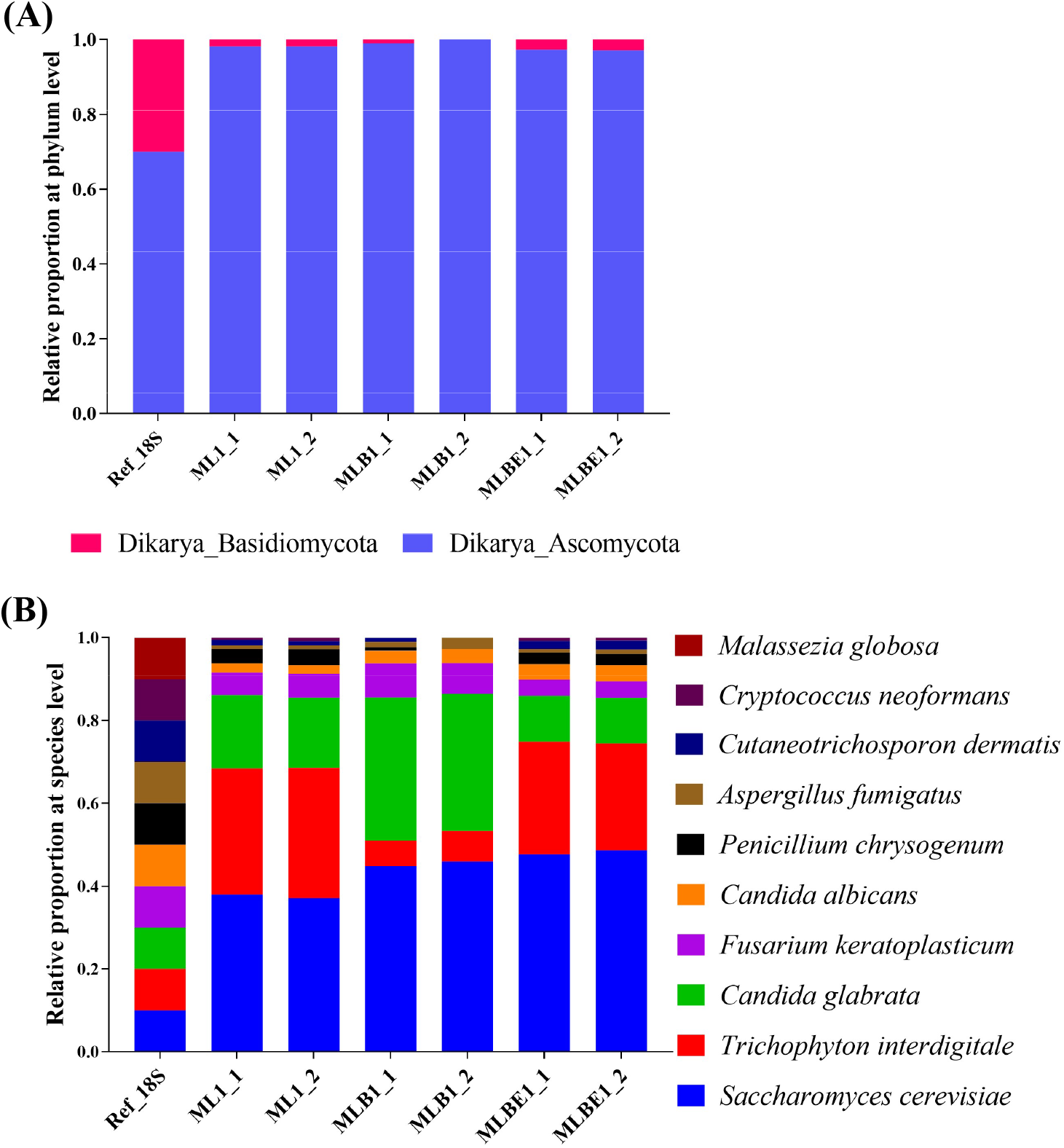
Relative bacterial abundance identified by Nanopore long-read sequencing using DNA isolated by 3 different extraction methods, lysis buffer alone (ML), incorporating bead-beating (MLB), or bead-beating plus MetaPolyzyme enzymatic treatment (MLBE). Data illustrated shows (A) all phyla, (B) species identified. The initial samples: ML1, MLB1, and MLBE1 were duplicated for Nanopore sequencing, abbreviated as ML1_1, ML1_2, MLB1_1, MLB1_2, MLBE1_1 and MLBE1_2, respectively. The reference data of 18S gene sequencing was abbreviated as Ref_18S.

At an individual phylum comparison, no significant difference between Basidiomycota and Ascomycota was found at ML and MLBE; however, a significant reduction of the Basidiomycota (*p =* 0.0412, Table S4) and elevation of Ascomycota were observed in MLB (*p =* 0.0412, Table S4) (Figure 6A).

**Figure 6:**
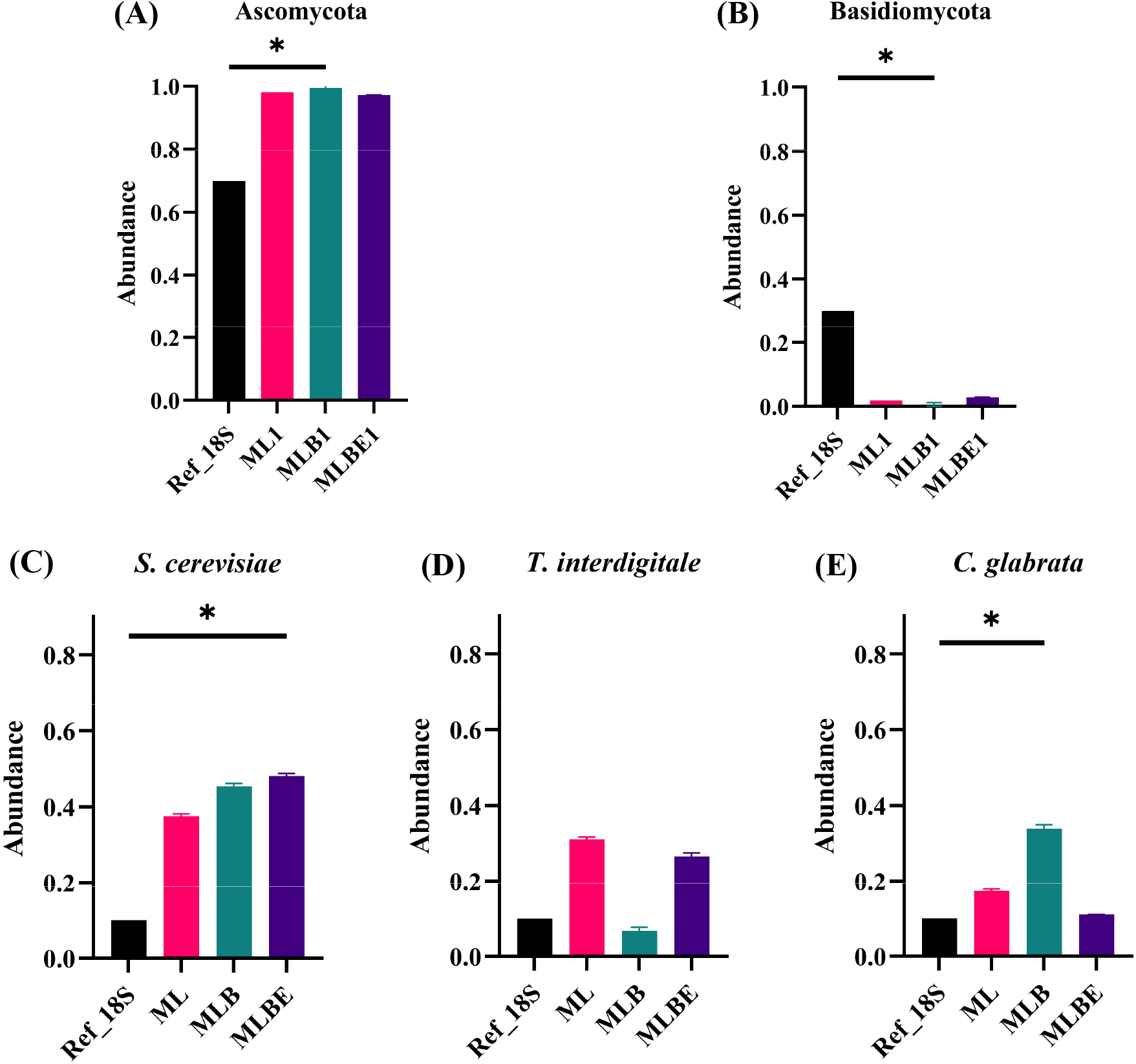
Relative abundance of top 3 fungal taxa identified by Nanopore sequencing of DNA isolated under different DNA extraction conditions; 96 MagBead DNA lysis buffer alone (ML), incorporating bead-beating (MLB), or bead-beating plus MetaPolyzyme enzymatic treatment (MLBE). After averaging duplicates, data illustrates the abundance of phyla (A) Ascomycota and (B) Basidiomycota; and species (C) *Saccharomyces cerevisiae*, (D) *Trichophyton interdigitale* and (E) *Candida glabrata*. Lines within the data plots indicate median values. Significant differences were observed, \**p* < 0.05 (Kruskal-Wallis test and Dunn’s multiple comparisons posthoc test).

At the genus level, the fungal proportions varied between the reference and the samples; Figure S5B. Comparing identified genera, MLB treatment significantly reduced *Penicillium*, *Cutaneotrichosporon,* and *Cryptococcus* (*p*-value of 0.0412, 0.0412, and 0.0395, respectively, Table S4); Figure S8. In addition, *Saccharomyces* was highly abundant in all treatment samples, with MLBE showing a significant increase compared to the reference data (*p* = 0.0412, Table S4). Furthermore, the large genus of filamentous fungi, *Fusarium*, was significantly decreased in abundance in the MLBE (*p* = 0.0412, Table S4) compared to ML and MLB. At the species level (Figure 6C-E); Figure S9, *Saccharomyces cerevisiae* and *Fusarium keratoplasticum* significantly differed between reference and MLBE (*p =* 0.0412, Table S4). Furthermore, *Candida glabrata* (*p =* 0.0412, Table S4), *Penicillium chrysogenum* (*p =* 0.0412, Table S4), *Cutaneotrichosporon dermatis* (*p =* 0.0412, Table S4) and *Cryptococcus neoformans* (*p =* 0.0395, Table S4) at MLB were varied in compared to the reference. At ML, *Candida albicans* are diverse compared to the reference data (*p =* 0.0412, Table S4); Figure S9.

Whilst there was no difference in the Chao1 index (Figure 7A), the Shannon diversity index showed a significant decrease in α-diversity in MLB (Figure 7B). For β-diversity analysis (Figure 7C-D), there was clear clustering based on treatment type. MLB showed the most similarity to the reference using this measure.

**Figure 7:**
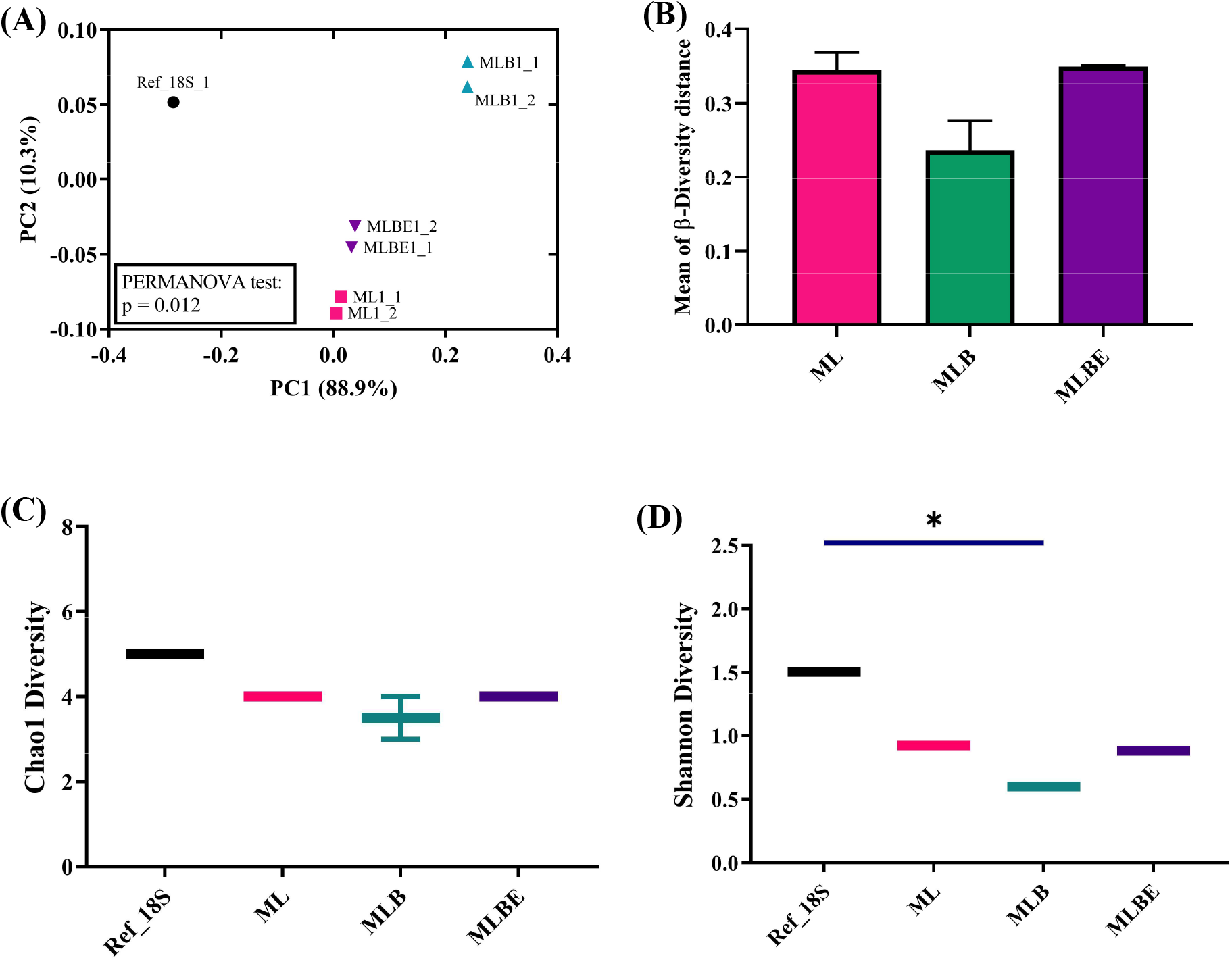
Comparison of fungal diversity following different extraction protocols. (A) β-diversity at the species level by Bray-Curtis metrics, (B) Coordinate distance of β-diversity at the fungal species level, (C) α-diversity at the species level using Chao1 index, (D) α-diversity at the species level using Shannon index. The bars of the histogram represent the mean with standard deviation. The line in the data plots indicates the median value. Significant differences were observed, \**p* < 0.05 (Kruskal-Wallis test and Dunn’s multiple comparisons posthoc test).

### The 18S rRNA primer pair can amplify all the mock community sample species

ClustalW multiple sequence alignment tool was utilised to evaluate the binding capabilities of the primer set (Fun18S1/FR-1), and it revealed the perfect alignment to all the fungal species in the mock microbial samples; Figure S1. Using the DNA extract from the known *M. globosa* colony, the same primer set was used to identify any amplification biases on the species from any lysis methods, proving no biases for the species. The result was confirmed as *M. globosa* using Nanopore and Sanger sequencing.

To ensure the feasibility of binding efficiency of the primer set in samples, all the samples of the same concentration were pooled. The pooled samples were spiked with DNA extract of known *M. globosa* at different concentrations: 50%, 25%, 10%, 5%, and 0% of *M. globosa*. Then, PCR was performed using the same primer set and proceeded to Nanopore sequencing. It revealed that the primer set could amplify the genus; however, it cannot be detected if the abundance is lower than 10%; Figure S2 and S3.

### DNA extraction provided imbalanced fungal proportion

To identify any DNA extraction bias, a specific primer set for the genus *Malassezia* (Mal1F and Mal1R) was utilised [50]. The amplified DNA bands on the gel are found in all samples along with multiple unspecific bands; therefore, we used the 1-in-3 dilution for the samples. In this experiment, ML2, MLB2, and MLBE1 samples were selected based on the band intensity to check the genus by Sanger sequencing. Using the BLAST alignment tool at the NCBI platform, the Sanger sequences were confirmed as *M. globosa*, showing the presence of the genus in the samples of all lysis conditions.

## DISCUSSION

Using bacterial and fungal mock samples, we compared the performance of three extraction protocols (with increasing treatment time and intensity) on the quantity and quality of DNA and the microbial abundance and diversity. To assess these lysis conditions, the bacterial standard data set was obtained from Zymo Research, at which 16S profiling was performed on the V3-V4 region of the ribosomal gene using the illumina^®^ MiSeq^TM^ (2×300 bp) platform [54]. For the assessment of fungal communities, proportional data was provided by ATCC, which assembled their genomes using Illumina and Oxford Nanopore Hybrid Assembly. As the current study utilised Oxford Nanopore sequencing technology, it may not be directly comparable; however, these platforms have previously been shown to be comparable in detecting microbial genera [55].

The standard dataset has 3 bacterial phyla: Firmicutes (83.29%), Bacteroidetes (12.47%) and Proteobacteria (4.24%). In the mock microbiota community samples, these phyla accounted for over 97% of the bacterial proportion, with the remaining 2-3% abundance attributed to Verrucomicrobia and Actinobacteria in all the conditions and Lentisphaerae only in MLBE1_2. Typically, Bacteroidetes (73.13 ± 22.16%), Firmicutes (22.2 ± 18.66%), Proteobacteria (2.15 ± 10.39%), Actinobacteria (1.82± 3%) and Verrucomicrobia (<1%) were the top 5 phyla found in healthy individuals [56, 57]. According to the online platform Integrated Microbial Genomes-Human Microbiome Project (IMG/HMP), Lentisphaerae, a member of the Planctomycetes-Verrucomicrobia-Chlamydia (PVC) superphylum, is one of the minor microbial phyla that can be identified in a healthy individual gut [58]. Since the reference kit was developed using faecal material from healthy adult donors [54], these phyla may originate from the original sample rather than a contaminant. Furthermore, 46 genera and 71 species were identified in the mock samples while the reference data reported 31 genera and only 5 species. These taxonomy data suggest that higher taxonomical resolution may be achieved with entire ribosomal gene sequencing by long-read Nanopore technology compared with short-read Illumina approaches [38, 39, 59]. Despite the limited information in the reference data, these techniques are comparable, as evidenced by Gehrig et al. [60], and therefore it was utilised as standard bacterial data to evaluate the lysis conditions.

The ATCC mycobiome reference data had 10 fungal species [61] from 9 genera and 2 phyla (70% relative abundance of Ascomycota and 30% of Basidiomycota). However, the mock samples were largely dominated by Ascomycota (∼98% relative abundance) and the remaining by the latter (Figure 5A). Therefore, the relative abundance of their downstream taxonomy, *Cryptococcus* and *Cutaneotrichosporon,* was very low (∼1%), and *Malassezia* was not identified in any lysis conditions (Figure 5B). The skew may be a result of extraction bias on the 18S rRNA gene or different sequencing techniques between reference and mock samples.

To ensure that the primers could amplify *M. globosa*, DNA was extracted from a pure culture and successfully amplified using the primer pair (MAL1F /MAL1R), suggesting that the primer set can amplify this species; Figure S2A. Subsequently, the mock microbiota community samples were amplified using the genus-specific primer set, and the genus was found, albeit a very proportion of *Malassezia*. We also evaluated the binding efficiency of the primer set on *Malassezia* in a matrix of samples by preparing spike-in controls. We found that the primers can bind and successfully amplify the *Malassezia* in a complex matrix when the DNA template contains at least 10% of it; Figure S2B and S3.

The genus *Malassezia* has been described as a difficult-to-lyse yeast [62]. Interestingly, the abundance decreased with increased bead beating in a previous study. It revealed that the relative abundance of *Malassezia* from no bead-beating to 15 min decreased from 2.8 to 0.1%, and the genus was not present after 20 min of bead-beating [63]. This suggests an enhanced sensitivity to bead-beating, which may be further exacerbated by the need for high-quality DNA for Nanopore sequencing. The detection of *Cryptococcus* was also inconsistent in MLB. This genus is also an encapsulated yeast, and it is difficult to isolate the DNA due to its thick and resistant capsule. Frau et al. identified that the DNA contents of *C. neoformans* were reduced following bead-beating [64]. In a study using an Illumina whole genome sequencing approach, 2 cycles of rapid bead-beating (45 s, 4.5 materials/sec by RiboLyser Homogenizer) were used to extract *C. neoformans* from the isolates [65]. Based on this, it may be assumed that longer bead beating time was to cause the reduction of the fungal species. These results show that optimising bead beating across different fungi is complex and optimal conditions may not be achievable for every type.

We also investigated the taxonomical differences in each lysis condition for each phylum, genus, and species using the 16S and 18S rRNA gene amplicons. A significant elevation in the relative abundance of the bacterial phylum Bacteroidetes (*p =* 0.0394, Table S3) was observed in the MLB condition, while the phyla in other states were not significantly different from the reference data (Figure 3A). At the genus level, the relative abundance of three common genera such as *Bacteroides* (*p =* 0.0412, Table S3)*, Clostridium IV* (*p =* 0.0412, Table S3), and *Anaerobacterium* (*p =* 0.0412, Table S3), was significantly elevated in MLB, and *Gemmiger* (*p =* 0.0412, Table S3) in MLBE was increased significantly. *Phascolarctobacterium* (*p =* 0.0412, Table S3) in ML was significantly decreased (Figure 3B). A study by Kumar et al. supported the findings that the relative abundance of *Bacteroides* and *Clostridioides* was increased in the method lacking bead beating process while *Phascolarctobacterium,* an abundant genus in the GI, had an elevated abundance with bead-beating [66]. In contrast, the genus *Phascolarctobacterium* was consistent across the lysis techniques (*p* > 0.05, Table S3), supporting that the bead-beating lysis could influence the recovery of the microbiota [66].

At the fungal phylum level (Figure 5A), 2 phyla were identified: Ascomycota and Basidiomycota. A reduction in the relative abundance of Basidiomycota and an increase in Ascomycota was observed across all the extraction and lysis conditions; however, only MLB reached a significant difference to reference data (*p =* 0.0412, Table S4). A significant decrease in the genus *Penicillium* (*p =* 0.0412, Table S4), *Cutaneotrichosporon* (*p =* 0.0412, Table S4), and *Cryptococcus* (*p =* 0.0395, Table S4) were observed using MLB. For the MLBE condition, *Saccharomyces* (*p =* 0.0412, Table S4) was increased, and *Fusarium* (*p =* 0.0412, Table S4) was decreased significantly. No significant differences in the genus were observed for the ML treatment compared to the reference data. Furthermore, no significant variation was found at the species level (Figure 5B). Considering the relative abundance of bacteria and fungi, ML and MLBE conditions yielded results most similar to the reference data. MLB treatment gave promising results for bacterial profiling but was not as promising for extracting high-quality fungal DNA.

The significant β-diversity (*p =* 0.033) of the bacterial species (Figure 4A) was found among different lysis conditions, and the PCoA distance analysis revealed that the ML and MLBE have the most similar distances to the reference (Figure 4B). No significant differences in bacterial α-diversity were identified (Figure 4C-D). Fungal β-diversity analysis was significantly different (*p =* 0.012) among all lysis techniques (Figure 7A), and ML and MLBE methods showed the same coordinate distance to reference (Figure 7B). No significant difference in α-diversity was found except for the MLB condition using the Shannon index (Figure 7D). Overall, it revealed that ML and MLBE have more similarities to the bacterial and fungal reference data than MLB.

This study revealed variation in DNA yield, which could influence downstream results. Therefore, higher sample numbers may provide more significant, more robust insights. Bead beating is considered critical for complete microbial lysis and accurate assessment of the relative abundance and diversity, particularly short-read V3-V4 amplicons [67]. However, full-length amplicon sequencing requires better DNA quality with less tolerance for sheared DNA. This, therefore, leads to a balance between maximal DNA yield using methods such as bead beating and reduced DNA quality for downstream amplification.

In the study, we combined existing primer sets for the full-length amplification of the 16S rRNA and 18S rRNA gene [28, 45] with an improved Phusion plus PCR protocol. For the 18S rRNA gene-specific primer, Banos et al. had previously evaluated the performance of the primer pair using an *in silico* analysis only [45]. The ClustalW Multiple Alignment via BioEdit tool revealed the alignment of the primers to all the mock species; Figure S1. Our study applied it specifically to amplify the gut mycobiome, which is largely recommended for further fungal amplicon sequencing studies.

In this study, we optimised the full-length 18S rRNA primer set (FUN18S1/FR-1). Considering the analytical performance, reagent cost, and processing time, ML treatment was identified as a straightforward, rapid, and efficient profiling method for bacterial and fungal profiles. This contrasts with many earlier studies that have highlighted the benefit of bead beating; however, we found that in the context of long read sequencing, the bead beating methodology had a negative impact on the amplification of full-length ribosomal genes compared to other methods. DNA shearing may not be problematic for short-read sequencing; however, it may limit the ability to amplify full-length amplicons for long-read sequencing.

The limitation of the study was that the sequencing techniques for the reference and mock samples were different, and therefore different levels of resolution were identified. There were potential technical biases during DNA extraction and sequencing processes, and all the profiling was based on a single extraction from each condition. Greater replication would yield more robust results. Despite its limitations, this is an initial proof-of-concept study prior to a gut microbiome study on human fecal samples.

To summarise, the study highlights the need to obtain high-quality DNA for profiling the human gut microbiome and mycobiome using long-read sequencing. This simple approach may also provide a time- and cost-efficient approach for simultaneously obtaining DNA to study bacteria and fungi residing in the gut.

## Acknowledgements

We extend our sincere thanks to the Center of Excellence in Systems Microbiology for enabling us to perform the study. The authors acknowledge the excellent technical support provided by Ms. Ariya Khamwut (Program in Medical Sciences; Faculty of Medicine; Chulalongkorn University) and also express their gratitude to Dr. Alessandra Frau and Dr. Rachael Slater (Institute of Systems Molecular and Integrative Biology; University of Liverpool) for the extraction and purification of DNA from spores of cultured *Penicillium chrysogenum* and *Malassezia globosa*.

## Ethics declarations

The authors declare that they have no conflicts of interest. This article contains no studies involving animals or human participants performed by the authors.

## Availability of data and materials

Data will be available as Supplementary Materials.

## Author contributions

B.J.C., J.L.F., M.S.T., N.H., P.C., S.P. and V.S. contributed to study conception and design; M.S.T., P.C. and V.S. performed experiments and analysed the data; V.S. and P.K. obtained the sequencing data, completed the database curation and the taxonomy files; B.J.C., J.L.F., N.H. and S.P. provided supervision, reviewed data and are the guarantors of the study; M.S.T. drafted the original manuscript; B.J.C., J.L.F., S.P. and N.H, contributed to the critical revision of the manuscript. All authors read and approved the final manuscript.

