## Supplementary Figures for "Optimisation of a DNA extraction protocol for improving the bacterial and fungal classification based on Nanopore sequencing"

### Supplementary Materials

##### Table S1: Primers used for PCR amplification of marker-specific genes

| Primers | *Oligonucleotide primer sequences (5′ to 3′) | Target Region | Target Gene | Reference |
|---|---|---|---|---|
| S-D-BACT-0008-C-S-20 (27F) | TTT CTG TTG GTG CTG ATA TTG CAG RGT TYG ATY MTG GCT CAG | V1-V9 | 16S rRNA | [1] |
| S-D-BACT-1492-A-A-22 (1492R) | ACT TGC CTG TCG CTC TAT CTT CCG GYT ACC TTG TTA CGA CTT | V1-V9 | 16S rRNA | [1] |
| NU-SSU-0068-5′-20 (FUN18S1) | TTT CTG TTG GTG CTG ATA TTG CCC ATG CAT GTC TAA GTW TAA | V1-V8 | 18S rRNA | [2] |
| NU-SSU-1648-3′ (FR-1) | ACT TGC CTG TCG CTC TAT CTT CAN CCA TTC AAT CGG TAN T | V1-V8 | 18S rRNA | [2] |

*Sequences underlined indicate the primer binding site

[1] Matsuo Y, Komiya S, Yasumizu Y, Yasuoka Y, Mizushima K, Takagi T, et al. Full-length 16S rRNA gene amplicon analysis of human gut microbiota using MinION nanopore sequencing confers species-level resolution. BMC Microbiol. 2021;21(1):35.

[2] Banos S, Lentendu G, Kopf A, Wubet T, Glöckner FO, Reich M. A comprehensive fungi-specific 18S rRNA gene sequence primer toolkit suited for diverse research issues and sequencing platforms. BMC Microbiology. 2018;18(1):190.

##### Table S2: Barcode adapters used for the preparation of DNA libraries

| *Adapters | Oligonucleotide primer sequences (5′ to 3′) |
|---|---|
| BC36/RB36 | ATGTCCCAGTTAGAGGAGGAAACA |
| BC37/RB37 | GCTTGCGATTGATGCTTAGTATCA |
| BC38/RB38 | ACCACAGGAGGACGATACAGAGAA |
| BC39/RB39 | CCACAGTGTCAACTAGAGCCTCTC |
| BC40/RB40 | TAGTTTGGATGACCAAGGATAGCC |
| BC41/RB41 | GGAGTTCGTCCAGAGAAGTACACG |
| BC42/RB42 | CTACGTGTAAGGCATACCTGCCAG |
| BC43/RB43 | CTTTCGTTGTTGACTCGACGGTAG |
| BC44/RB44 | AGTAGAAAGGGTTCCTTCCCACTC |
| BC45/RB45 | GATCCAACAGAGATGCCTTCAGTG |
| BC46/RB46 | GCTGTGTTCCACTTCATTCTCCTG |
| BC47/RB47 | GTGCAACTTTCCCACAGGTAGTTC |
| BC48/RB48 | CATCTGGAACGTGGTACACCTGTA |

*The adapters were obtained from PCR Barcoding Expansion 1-96 (EXP-PBC096) kit (Oxford Nanopore Technologies, UK).

**Table S3: A table of *p*-values for the comparison of different lysis conditions at the phylum, genus and species level of bacteria**

| Classification | *p*-values* | | |
|---|---|---|---|
| | **Ref vs ML** | **Ref vs MLB** | **Ref vs MLBE** |
| **Phylum** | | | |
| Firmicutes | ns | ns | ns |
| Bacteroides | ns | 0.0429 | ns |
| Proteobacteria | ns | ns | ns |
| **Genus** | | | |
| *Faecalibacterium* | ns | ns | ns |
| *Bacteroides* | ns | 0.0412 | ns |
| *Lachnospiracea* | ns | ns | ns |
| *Clostridium IV* | ns | 0.0412 | ns |
| *Gemmiger* | ns | ns | 0.0412 |
| *Anaerobacterium* | ns | 0.0412 | ns |
| *Blautia* | ns | ns | ns |
| *Alistipes* | ns | ns | ns |
| *Roseburia* | ns | ns | ns |
| *Ruminococcus* | ns | ns | ns |
| *Parabacteroides* | ns | ns | ns |
| *Phascolarctobacterium* | 0.0412 | ns | ns |
| *Fusicatenibacter* | ns | ns | ns |
| **Species** | | | |
| *Faecalibacterium sp.* | ns | ns | ns |
| *Lachnospiracea sp.* | ns | ns | ns |
| *Bacteroides sp.* | ns | ns | ns |
| *Gemmiger sp.* | ns | ns | 0.0412 |
| *Clostridium IV sp.* | ns | ns | ns |
| *Bacteroides vulgatus* | ns | ns | ns |
| *Anaerobacterium sp.* | ns | 0.0412 | ns |
| *Ruminococcus sp.* | ns | ns | ns |
| *Phascolarctobacterium sp.* | 0.0412 | ns | ns |
| *Roseburia sp.* | ns | ns | ns |
| *Fusicatenibacter sp.* | ns | ns | ns |
| *Parabacteroides sp.* | ns | ns | ns |
| Megamonas sp. | ns | ns | ns |
| Bacteroides dorei | ns | ns | ns |
| Vampirovibrio sp. | ns | ns | ns |

*The *p*-values were obtained from Dunn's multiple comparison test. ns, not significant ($p > 0.05$).

Optimisation of DNA extraction protocol for improving the bacterial and fungal classification based on Nanopore sequencing. Thu MS, Sawaswong V, Chanchaem P, Campbell BJ, Hirankarn N, Fothergill JL, Payungporn S.

**Table S4: A table of *p*-values for the comparison of different lysis conditions at the phylum, genus and species level of fungi**

| Classification | *p*-values* | | |
|---|---|---|---|
| | **Ref vs ML** | **Ref vs MLB** | **Ref vs MLBE** |
| **Phylum** | | | |
| Basidiomycota | ns | 0.0412 | ns |
| Ascomycota | ns | 0.0412 | ns |
| **Genus** | | | |
| *Saccharomyces* | ns | ns | 0.0412 |
| *Trichophyton* | ns | ns | ns |
| *Candida* | ns | ns | ns |
| *Fusarium* | ns | ns | 0.0412 |
| *Penicillium* | ns | 0.0412 | ns |
| *Aspergillus* | ns | ns | ns |
| *Cutaneotrichosporon* | ns | 0.0412 | ns |
| *Cryptococcus* | ns | 0.0395 | ns |
| **Species** | | | |
| *Saccharomyces cerevisiae* | ns | ns | 0.0412 |
| *Trichophyton interdigitale* | ns | ns | ns |
| *Candida glabrata* | ns | 0.0412 | ns |
| *Fusarium keratoplasticum* | ns | ns | 0.0412 |
| *Candida albicans* | 0.0412 | ns | ns |
| *Penicillium chrysogenum* | ns | 0.0412 | ns |
| *Aspergillus fumigatus* | ns | ns | ns |
| *Cutaneotrichosporon dermatis* | ns | 0.0412 | ns |
| *Cryptococcus neoformans* | ns | 0.0395 | ns |

*The *p*-values were obtained from Dunn's multiple comparison test. ns, not significant ($p > 0.05$).

**Figure S1**. **CLUSTALW Multiple sequence alignment.** Multiple alignment of the 18S rRNA gene sequences in mock microbiota community, aligned to **A)** forward and **B)** reverse oligonucleotide primers using CLUSTALW (www.genome.jp/tools-bin/clustalw ; accessed 13 June 2022). Full sequences of 7 species were found from SILVA database (www.arb-silva.de/; accessed 13 June 2022). For *Malassezia*, the full 18S rRNA was retrieved using Nucleotide Basic Local Alignment Search Tool (BLASTn). For *Trichophytum* and *Fusarium* genera, pair-alignment was performed using the contigs from ATCC genome assembly, then made a BLASTn at NCBI (https://blast.ncbi.nlm.nih.gov/Blast.cgi ; accessed on 13 June 2022).

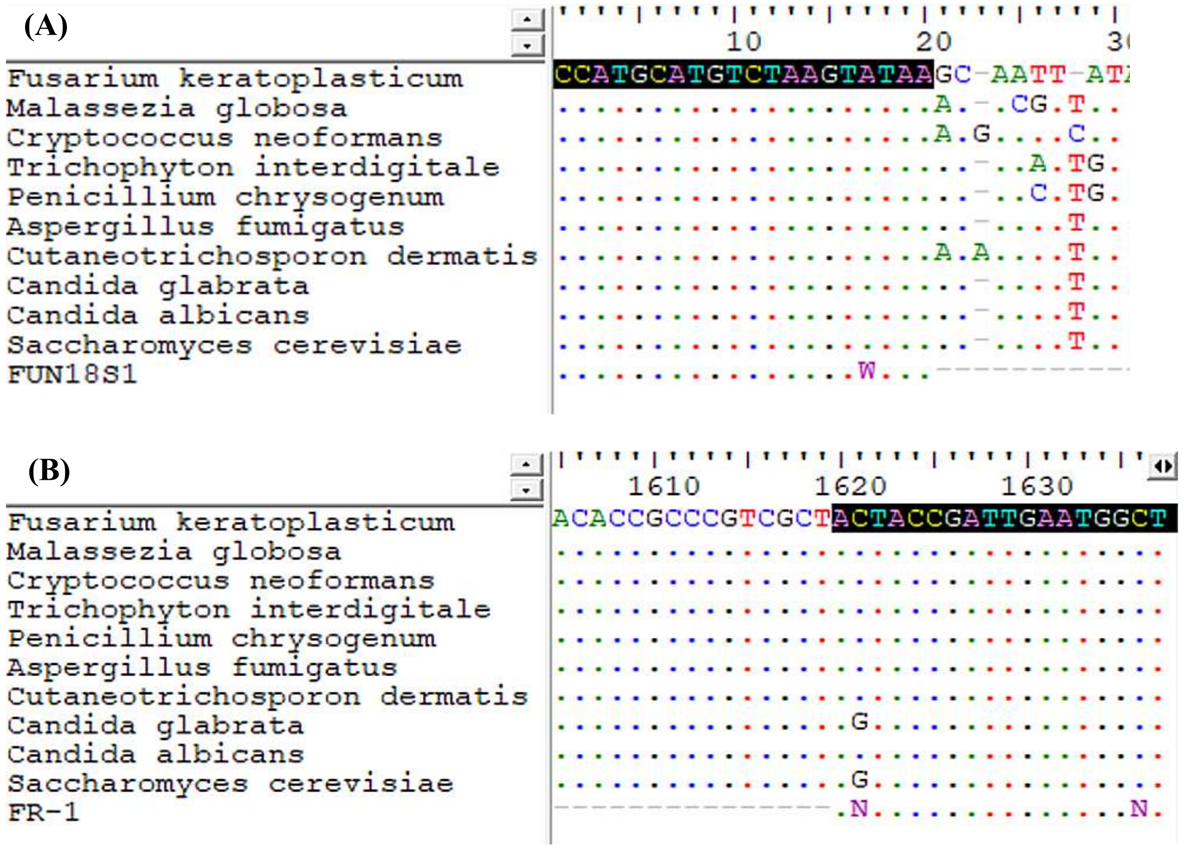

**Methods S1: Culture of, and isolation of DNA from, *Malassezia globosa.*** *M. globosa* was grown initially on modified Dixon's (mDixon) agar containing 26 mg/L chloramphenicol for 3–4 d, at 32 °C, as per conditions previously described for *Malassezia* sp. [Frau et al., 2019]. Briefly, mDixon broth was then inoculated with a single colony and incubated at 32 °C, shaking at 200 rpm for 48 h. Spores were harvested, and counted with a Neubauer-improved haemocytometer. And then extracted using a QIAamp Fast DNA mini kit (Qiagen; Hilden, Germany). Following extraction, DNA was quantified with a Qubit (Qubit dsDNA HS assay Kit; Life Technologies).

Frau, A., Kenny, J.G., Lenzi, L. *et al.* DNA extraction and amplicon production strategies deeply influence the outcome of gut mycobiome studies. *Sci Rep* 2019; **9**, 9328

**Figure S2. PCR amplification of *Malassezia* from isolated DNA extracted under different lysis conditions.** Gel electrophoresis of PCR amplicons generated using, A) the first PCR product in 2-step amplification to evaluate the amplification bias of the primer set on *Malassezia globosa* (MG) at which a negative control was performed but not shown here; B) using 18S rRNA primer set (FUN18S1/FR-1) on samples spiked with known amounts of purified *M. globosa* DNA. Different spike-in concentrations: 50%, 25%, 10%, 5%, and 0% of the mock community DNA library, were prepared using the same concentration of a mixture of mock control and known *M. globosa* samples; and C) using *Malassezia*-specific PCR primers MAL1F (5'-TCTTTGAACGCACCTTGC-3') and MAL1R (5'- AHAGCAAATGACGTATCATG-3') on all samples prior to Sanger sequencing. Positive (Pos) control, with 2.35 ng/µL DNA isolate from *M. globosa.* Negative control (Neg), with no DNA template included in the amplification.

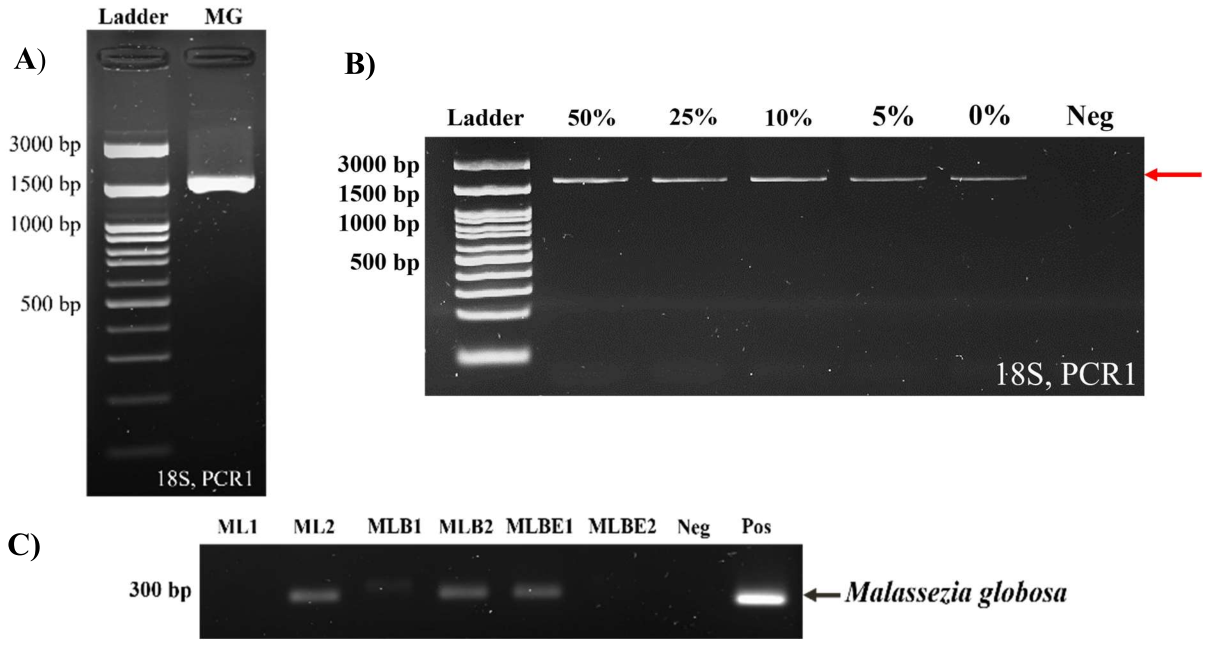

**Figure S3: Comparison of fungal abundance at spike-in controls.** Different spike-in concentrations: 50%, 25%, 10%, 5%, and 0%, were prepared using the same concentration of a mixture of mock microbiota community and known proportions of *M. globosa* samples.

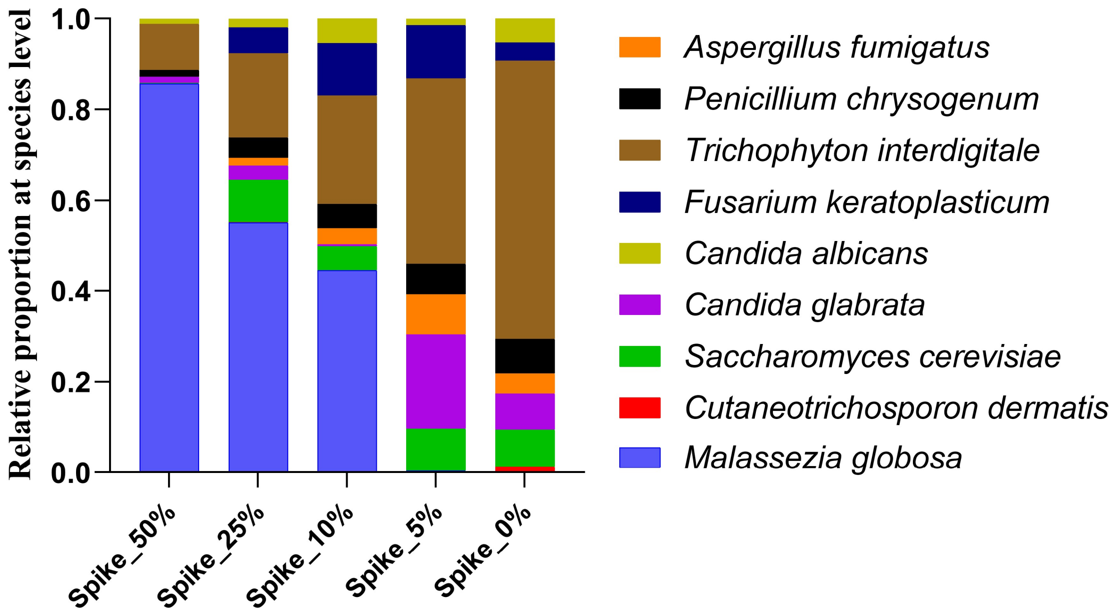

**Figure S4:** Rarefaction curves showing observed species richness based on (**A**) 16S rRNA, and (**B**) 18S rRNA gene sequencing, created by MicrobiomeAnalyst (www.microbiomeanalyst.ca/, accessed 04 June 2022). Three sample lysis conditions were applied to a mock microbial community including known bacterial and fungal species, using the 96 MagBead DNA lysis buffer (ML) alone, incorporating bead-beating (MLB), or bead-beating following MetaPolyzyme enzymatic pre-treatment (MLBE). The initial samples: ML1, MLB1, and MLBE1 were duplicated for Nanopore sequencing and abbreviated as ML1_1, ML1_2, MLB1_1, MLB1_2, MLBE1_1 and MLBE1_2. The 16S and 18S rRNA gene sequencing reference data were abbreviated as Ref_16S and Ref_18S, respectively. Species richness is the count of each operational taxonomy unit (OTUs) in each reference (Ref_16S and Ref_18S) and sample. The sequence sample size is the total read count of each reference (Ref_16S and Ref_18S) and sample.

**(A)**

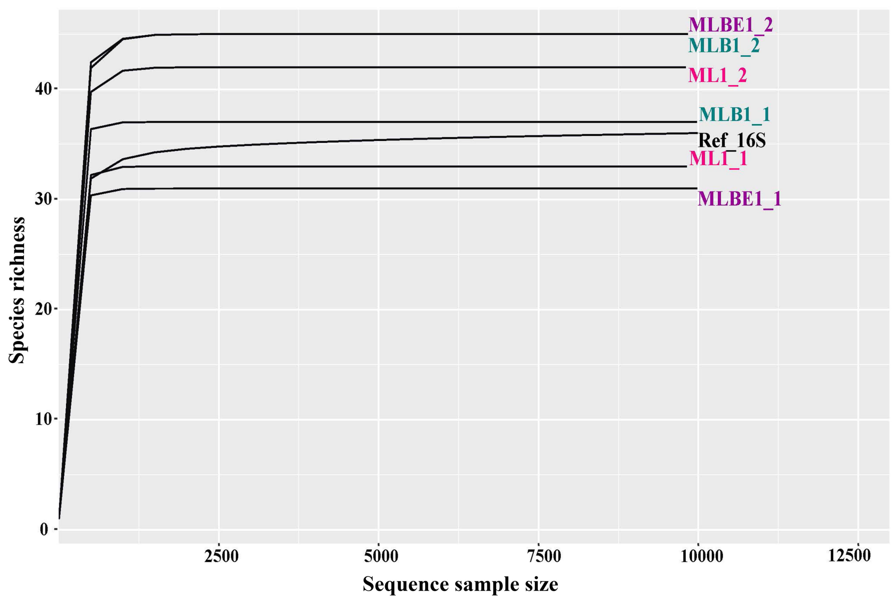


**(B)**

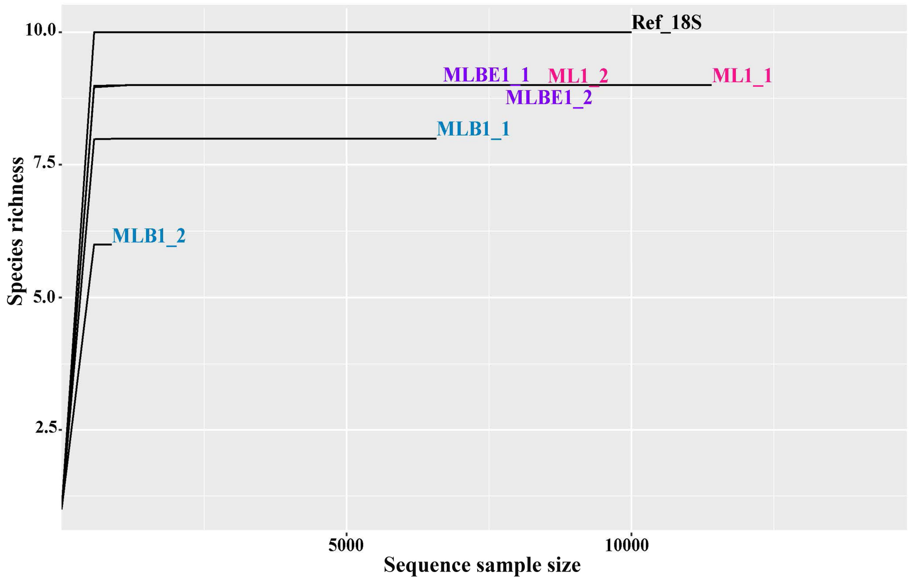

**Figure S5: Relative abundance identified by Nanopore long-read sequencing using DNA isolated by 3 different extraction methods**, lysis buffer alone (ML), incorporating bead-beating (MLB), or bead-beating plus MetaPolyzyme enzymatic treatment (MLBE). Data illustrated shows (A) the top 15 bacterial genera, and (B) all the fungal genera identified. The initial samples: ML1, MLB1, and MLBE1 were duplicated for Nanopore sequencing, abbreviated as ML1_1, ML1_2, MLB1_1, MLB1_2, MLBE1_1 and MLBE1_2, respectively. The reference data of 16S and 18S rRNA gene sequencing was abbreviated as Ref_16S and Ref_18S, respectively.

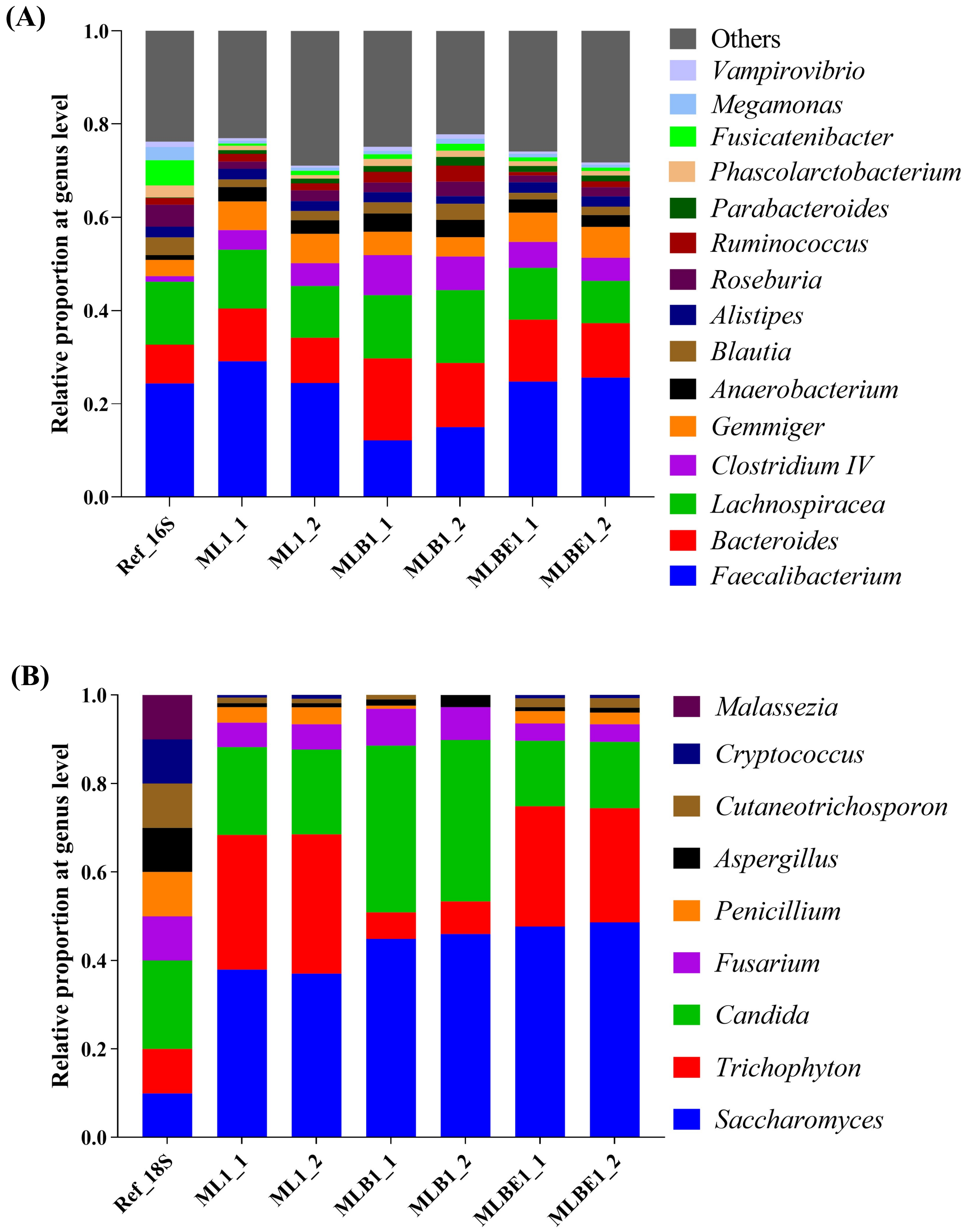

**Figure S6: Relative abundance of Top 15 bacterial taxonomy at the genus level.** Kruskal-Wallis statistic test; asterisks indicate adjusted *p*-value < 0.05 in Dunn's multiple comparisons posthoc test. Error bars are indicated on top of the boxes.

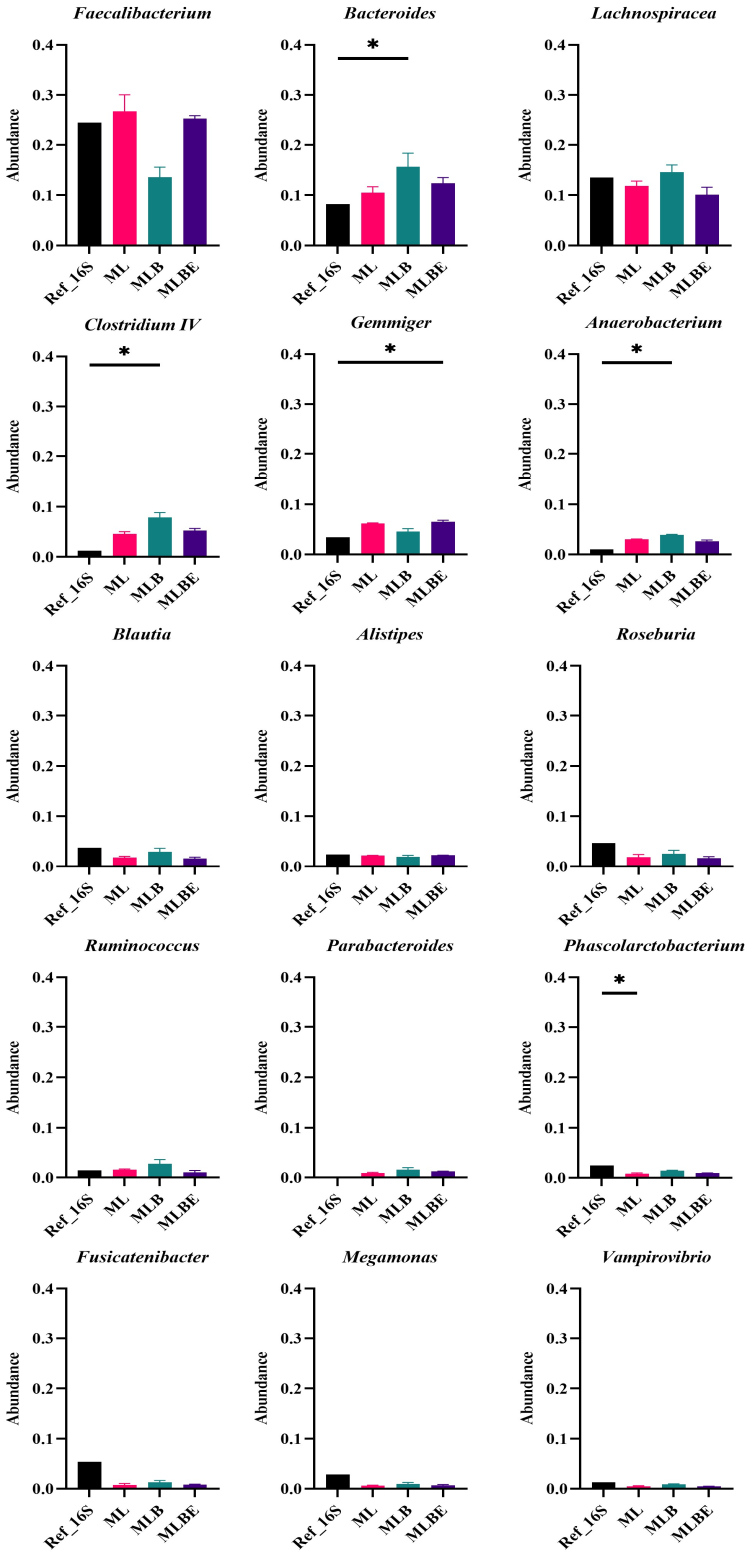

**Figure S7. Relative abundance of Top 15 bacterial taxa at the species level.** Kruskal-Wallis statistic test; asterisks indicate adjusted *p*-value < 0.05 in Dunn's multiple comparisons posthoc test. Error bars are indicated on top of the boxes.

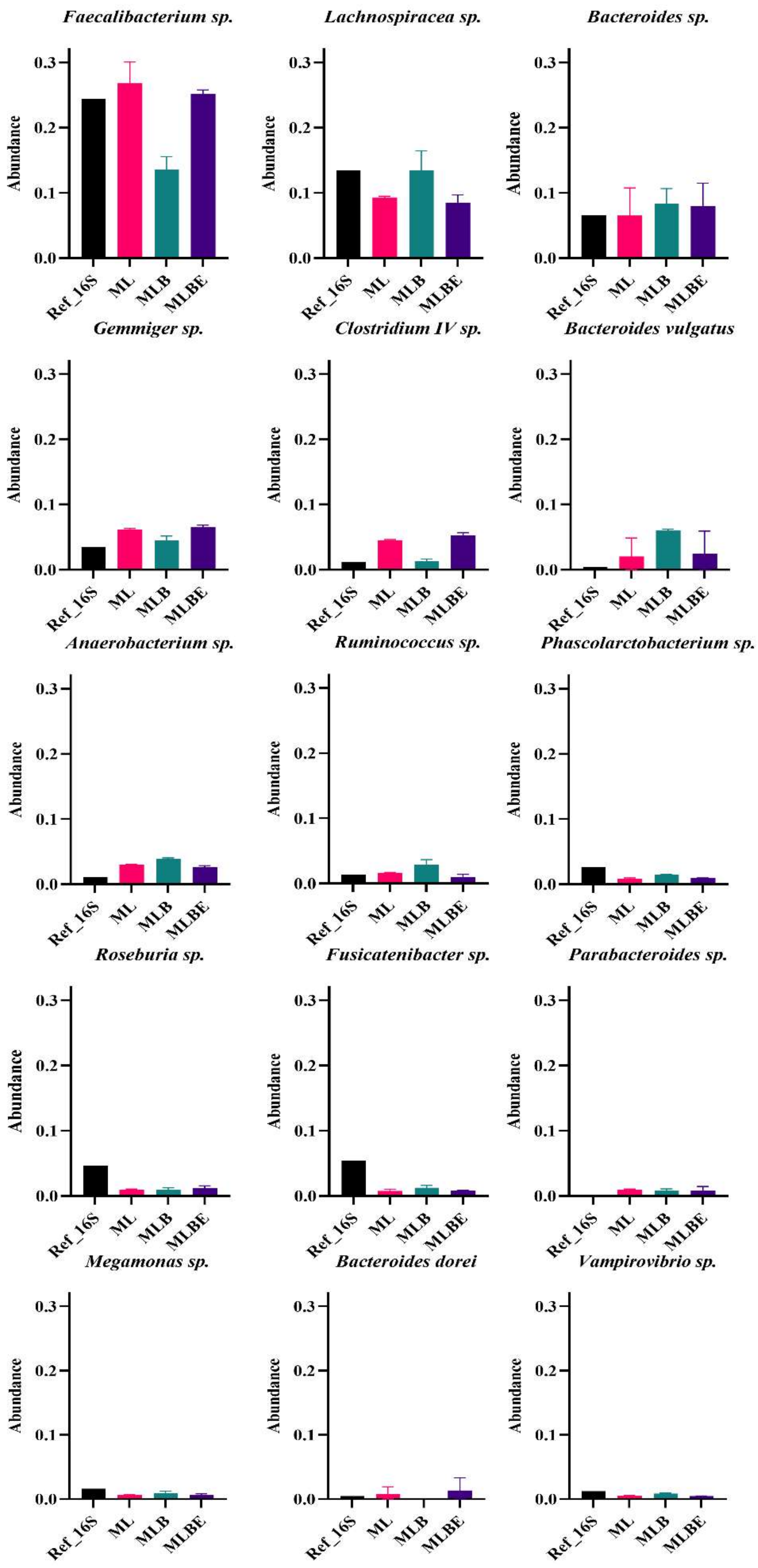

**Figure S8. Relative abundance of all fungal taxa identified at the genus level.** Kruskal-Wallis statistic test; asterisks indicate adjusted *p*-value < 0.05 in Dunn's multiple comparisons posthoc test. Error bars are indicated on top of the boxes.

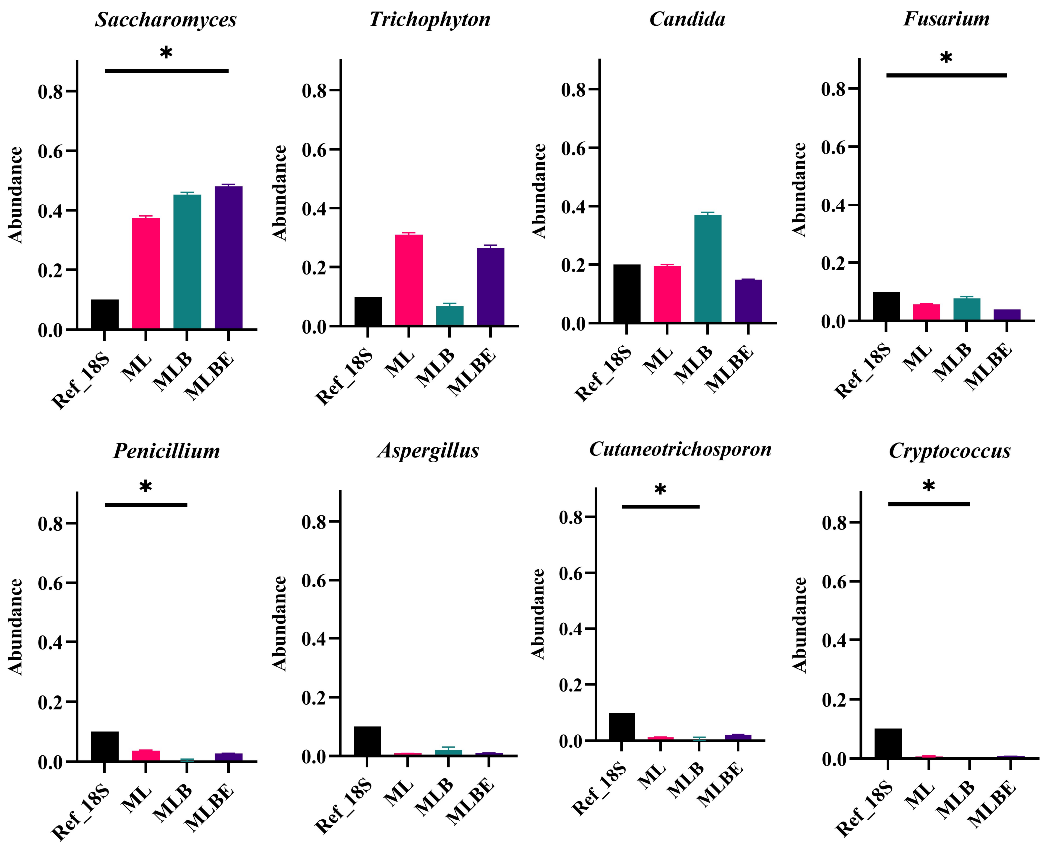

**Figure S9. Relative abundance of all fungal taxa identified at the species level.** Kruskal-Wallis test; asterisks indicate adjusted $p$-value < 0.05 in Dunn's multiple comparisons posthoc test. Error bars are indicated on top of the boxes.

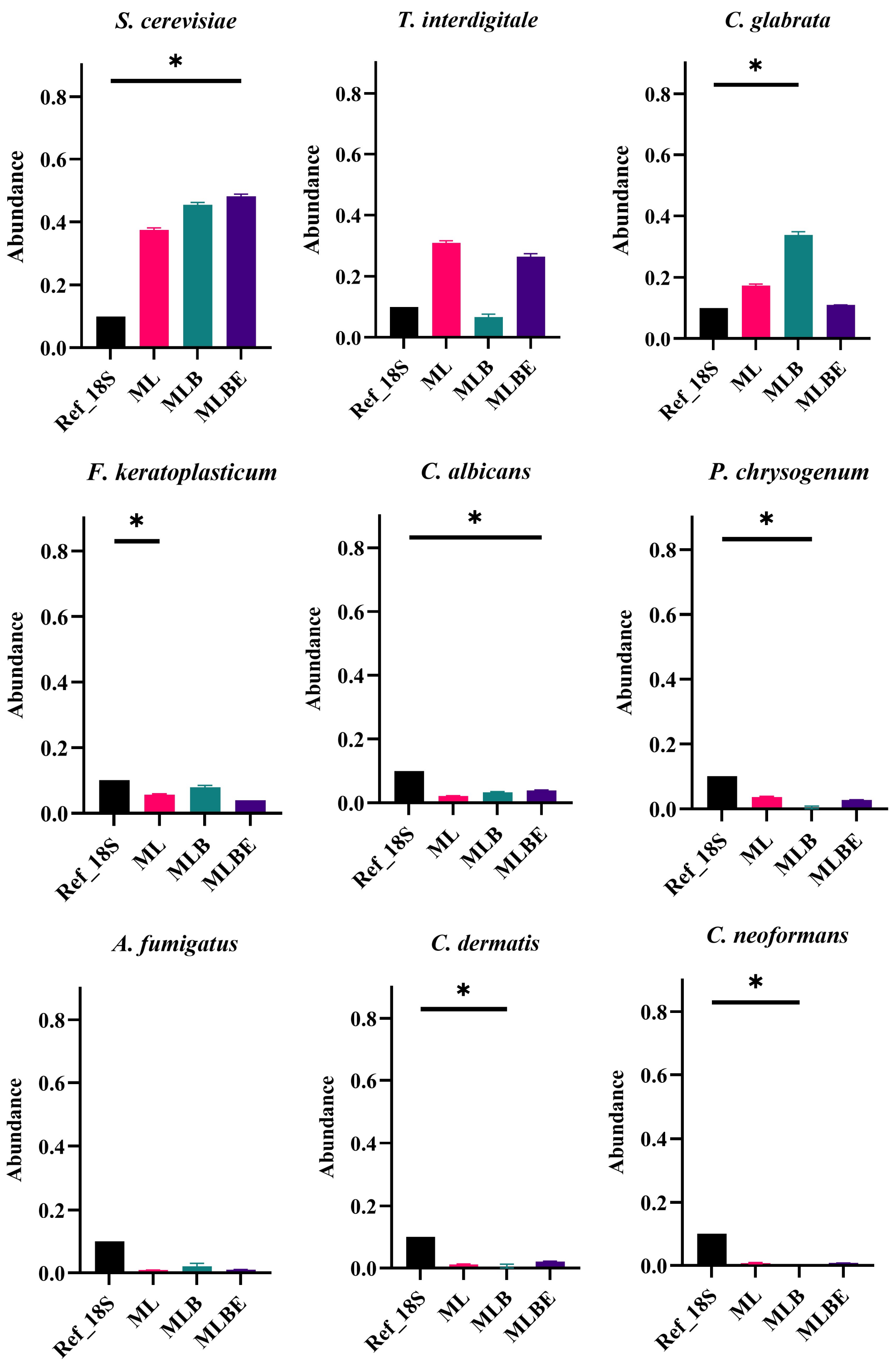
